# Identification of anti-fungal bioactive terpenoids from the bioenergy crop switchgrass (*Panicum virgatum*)

**DOI:** 10.1101/2023.02.24.529965

**Authors:** Xingxing Li, Ming-Yi Chou, Gregory M. Bonito, Robert L. Last

**Author notes:** Xingxing Li and Ming-Yi Chou contributed equally. Author for contact Robert Last, Department of Biochemistry and Molecular Biology, Michigan State University, 603 Wilson RD, East Lansing, MI 48823 USA. Department of Plant Pathology, University of Wisconsin-Madison, Madison, WI, 53706 USA.

## Abstract

Plant derived bioactive small molecules have attracted attention of scientists across fundamental and applied scientific disciplines. We seek to understand the influence of these phytochemicals on functional phytobiomes. Increased knowledge of specialized metabolite bioactivities could inform strategies for sustainable crop production. Our recent investigations of metabolomes of the upland and lowland ecotypes of the bioenergy crop switchgrass (*Panicum virgatum*) revealed large differences in types and abundances of terpenoid specialized metabolites. We hypothesized that – consistent with accumulating evidence that switchgrass genotype impacts microbiome assembly – differential terpenoid accumulation contributes to switchgrass ecotype-specific microbiome composition. An initial *in vitro* plate-based disc diffusion screen of 18 switchgrass root derived fungal isolates revealed differential responses to upland- and lowland-isolated metabolites. To identify specific fungal growth-modulating metabolites, we tested fractions from root extracts on three ecologically important fungal isolates – *Linnemania elongata, Trichoderma* sp. and *Fusarium* sp. Saponins and diterpenoids were identified as the most prominent antifungal metabolites. Finally, analysis of liquid chromatography-purified terpenoids revealed fungal inhibition structure – activity relationships (SAR). Saponin antifungal activity was primarily determined by the number of sugar moieties – saponins glycosylated at a single core position were inhibitory whereas saponins glycosylated at two core positions were inactive. Saponin core hydroxylation and acetylation were also associated with reduced activity. Diterpenoid activity required the presence of an intact furan ring for strong fungal growth inhibition. This study demonstrates that switchgrass genotype-specific metabolites differentially inhibit fungal isolates from the root and rhizosphere, supporting the hypothesis that these small molecules contribute to microbiome assembly and function.

## Introduction

Renewable energy produced from plant biomass grown with ecologically sound practices can contribute to a more secure, environmentally sustainable and economically stable future (Langholtz, Stokes and Eaton, 2016). Switchgrass (*Panicum virgatum*; Poaceae) has attracted attention as a biofuel feedstock because it is a perennial C4 photosynthetic plant with high cellulosic content, which thrives on land of little or no agricultural value (Sanderson et al., 2006; Casler et al., 2015). Switchgrass also exhibits substantial resistance to pests and diseases (Uppalapati et al., 2013; Casler et al., 2015). As climate change is predicted to cause shifts in pest and microbial disease pressure that would endanger biomass production (Jägermeyr *et al*., 2021), there is increasing opportunity to breed or metabolically engineer disease resistant varieties. Unlike domesticated crops, switchgrass retains tremendous phenotypic diversity, including in plant protective specialized metabolites (Casler et al., 2015; Lowry et al., 2019; Li et al., 2022).

The interactions of plants with pathogenic and beneficial microbes are determined by many factors, including production of specialized metabolites, whose synthesis and structures vary across the plant kingdom (Maeda and Fernie, 2021). Terpenoids are an excellent example as they are the largest characterized plant natural product family (Pichersky and Raguso, 2018). The protective roles of terpenoid specialized metabolites have been extensively explored in the model plant, *Arabidopsis thaliana*, and major Poaceae crops. These include directly inhibiting microbial pathogen growth as toxins and priming plant defense. For example, the nonvolatile avenacin triterpenoid saponins protect oats from the fungal pathogen, *Gaeumannomyces tritici*, which causes ‘take-all’ disease (Burkhardt et al., 1964; Bowyer et al., 1995; Papadopoulou et al., 1999). In addition, the labdane diterpenoids that are broadly found in rice, wheat and maize have demonstrated defensive roles as antimicrobial phytoalexins (Schmelz *et al*., 2014; Mafu *et al*., 2018; Ding *et al*., 2019; Murphy and Zerbe, 2020). Finally, C10 monoterpene and C15 sesquiterpene volatiles induce systemic acquired resistance within and between plants in *Arabidopsis* (Riedlmeier et al., 2017; Frank et al., 2021).

The microbiome has demonstrable impact on plant growth and can act as an extension of the plant immune system (Trivedi *et al*., 2020). Plants can recruit specific strains of microbes to help defend pathogen through plant-microbe coevolution. At a community level, studies showed plant associated microbiome composition plays a crucial role in determining disease incidence and severity (Wei et al., 2019; Chou et al., 2021). Manipulation of microbiomes was demonstrated to be a viable approach for plant disease control via microbe-mediated pathogen suppression (Fierer, 2017) or plant immune system priming (Raaijmakers and Mazzola, 2016). Rhizosphere microbial compositions and functions are influenced by multiple factors, such as abiotic conditions (Edwards *et al*., 2015; Simonin *et al*., 2020), plant species (Berg and Smalla, 2009; Fitzpatrick *et al*., 2018) and plant genotypes (Wagner et al., 2016; Morella et al., 2020; Brown et al., 2021; Beschoren da Costa et al., 2022). Among these, variations in metabolites of plant above- and below-ground tissues as well as root exudates have been implicated in microbiome function (Jones et al., 2009; Venturi & Keel, 2016; Korenblum et al., 2020; VanWallendael et al., 2022).

Species- or genotype-specific specialized metabolites play roles in mediating rhizosphere microbial interactions. For example, secretion of coumarin and benzoxazinoid specialized metabolites from the root shape the maize rhizosphere microbiomes (Stringlis et al., 2018; Cotton et al., 2019). There is increasing evidence suggesting that switchgrass genotype influences the rhizosphere microbiome (Beschoren da Costa *et al*., 2022; Ulbrich *et al*., 2022). Our working hypothesis is that genotype-specific differences in specialized metabolite accumulation influences rhizosphere microbial assembly.

As part of a larger effort to develop switchgrass varieties with high yield and low environmental impacts, we are engaged in studies of switchgrass metabolome screening by liquid chromatography–mass spectrometry (LC–MS) and other analytical chemistry methods. We previously identified close to 100 terpenoid metabolites, including saponins, diterpenes and sesquiterpenes, in the northern upland and southern lowland switchgrass ecotypes (Li et al., 2022; Tiedge et al., 2022). We demonstrated specific differences in metabolites across the terpenoid classes driven by ecotype, tissue/developmental stage and soil moisture, raising questions about their possible biological functions. Filling this knowledge gap will help lead to rational breeding of highly productive bioenergy crops with improved microbiome traits.

Here, we focused on abundant terpenoids that are dramatically different in root metabolomes among switchgrass ecotypes. We asked whether fungi isolated from switchgrass microbiomes are differentially sensitive to these terpenoids. We first showed that the upland and lowland switchgrass root extracts differentially inhibited the growths of 18 switchgrass rhizosphere fungi. Fungal bioactivity-guided metabolite fractionation (**Fig 1A**) revealed that fractions containing saponins and diterpenoids inhibit fungal growth. The hypothesis was tested and confirmed through fungal bioassays using a set of high-performance liquid chromatography (HPLC) fractions containing specific steroidal saponins and diterpenoids. Together, our results revealed that terpenoids are active against diverse fungi in the switchgrass root microbiome and indicate that these bioactive natural products are candidates as mediators of switchgrass ecotype-specific microbiome recruitment.

**Figure 1.**
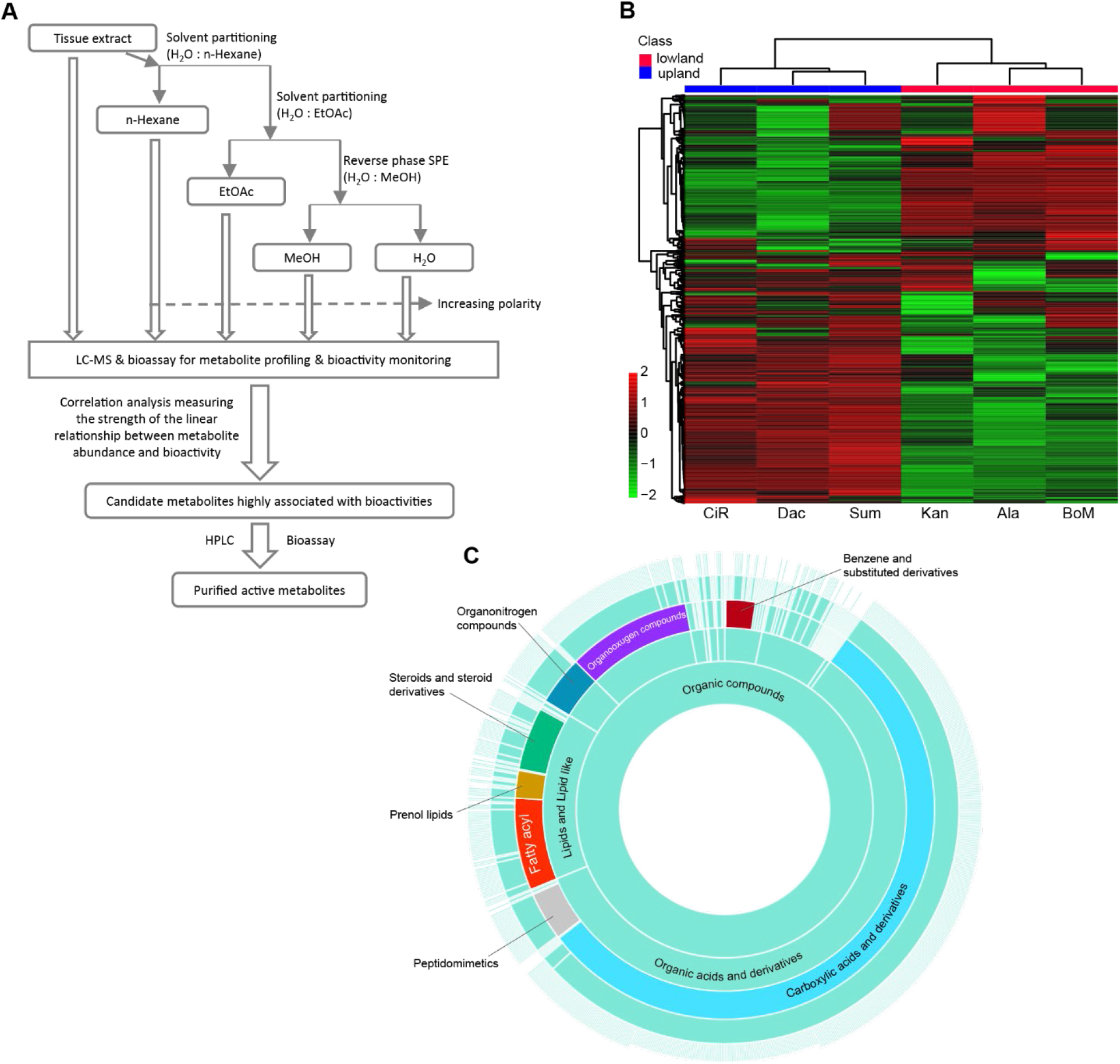
Workflow for the bioassay-guided identification of bioactive switchgrass metabolites and global view of switchgrass root tissue metabolomes revealed by untargeted metabolomics. (A) A workflow for isolation of bioactive compounds from switchgrass modified from Figure 2 from Ebada et al. (2008). (B) Heatmap generated using Euclidean correlation-based clustering (algorithm: Ward) of Log10 Fold Change (FC) of the 1777 metabolite features, identified in the root extracts of six switchgrass cultivars, with normalized abundance ≥ 500 arbitrary unit. Heatmap scale shows FC values. (C) Sunburst plot showing the CANOPUS classifications (Dührkop et al., 2021) for switchgrass metabolites with *m/z* values ≤ 850 based on MS/MS spectra of the metabolite features. Inner rings represent parental classes of outer rings (from inside to outside: kingdom – super class – class – subclass – level5). On the compound class level, the eight largest classes are colored and labeled.

## Results

### Terpenoids are ecotype-specific root metabolites

We followed up on previous observations that switchgrass root specialized metabolite profiles differ between upland and lowland ecotypes. We reasoned that a better annotated switchgrass metabolome would help to identify correlations between metabolites and antifungal bioactivities of root extracts from different switchgrass ecotypes/genotypes (Li *et al*., 2022). After extracting metabolites from large amounts of growth chamber grown root tissue biomass from the three upland and three lowland switchgrass cultivars (**Materials and Methods**), we evaluated the metabolites by LC-MS. Euclidean correlation-based clustering separated the three upland and lowland cultivars, confirming that these extracts have dramatically different metabolite constituents (**Fig 1B, Table S1**). Analysis of this untargeted metabolomics dataset using CANOPUS (class assignment and ontology prediction using mass spectrometry), a computational method for systematic compound class annotation (Dührkop *et al*., 2021), assigned chemical classes to the features with *m/z* values ≤ 850 (**Materials and Methods, Table S2**). This yielded a better understanding of the chemical composition of switchgrass root tissues, compared to annotating the features based on their relative mass defect (Li *et al*., 2022). Approximately half of the annotated features were carboxylic acids and derivatives, ∼14% lipids and lipid derived metabolites and ∼10% organooxygen compounds, which are mostly carbohydrates and carbohydrate conjugates (**Fig 1C, Table S2**).

To identify the major contributing metabolite classes to the divergent switchgrass root metabolomes, we fractioned the extracts by polarity using two phase separations with water/hexane followed by water/ethyl acetate (EtOAc) and then C18 resin solid phase extraction (SPE) eluted with water/methanol (**Fig 1A**). The metabolite fold-change revealed that the medium- and low-polarity metabolites contribute to the overall profile divergence more than do those small molecules of high-polarity (**Fig 2A**). Next we performed Orthogonal Projections to Latent Structures Discriminant Analysis (OPLS-DA, Wiklund et al., 2008) to identify the ecotype-specific differentially accumulating features (DAFs) that drive the metabolite profile separation in each fraction (**Fig 2B, Fig S1 A – D**). The analysis generated an S-plot from which the DAFs with high reliability and magnitude were selected according to a defined threshold (**Fig 2C, Fig S1 E – H**): 241 and 129 of the most highly weighted DAFs were identified for upland and lowland ecotypes, respectively (**Fig 2D, Table S 3 & 4**). Using a combination of CANOPUS, LC-MS databases and manual annotation (**Material and Methods**), we classified 39% and 43% of the upland and lowland DAFs into 10 chemical categories (**Fig 2D, Table S 3 & 4**). Notably, in the upland ecotype, 12% of DAFs were classified as C20 di-, C15 sesqui-, and C10 monoterpenoids, followed by 9% of DAFs grouped into the saponin class (C27-C30 core glycosylated triterpenoids and steroids) (**Fig 2D, Table S 3 & 4**). In strong contrast, 24% of lowland DAFs were classified as saponins while none were annotated as di-, sesqui- or monoterpenoids (**Fig 2D, Table S 3 & 4**).

**Figure 2.**
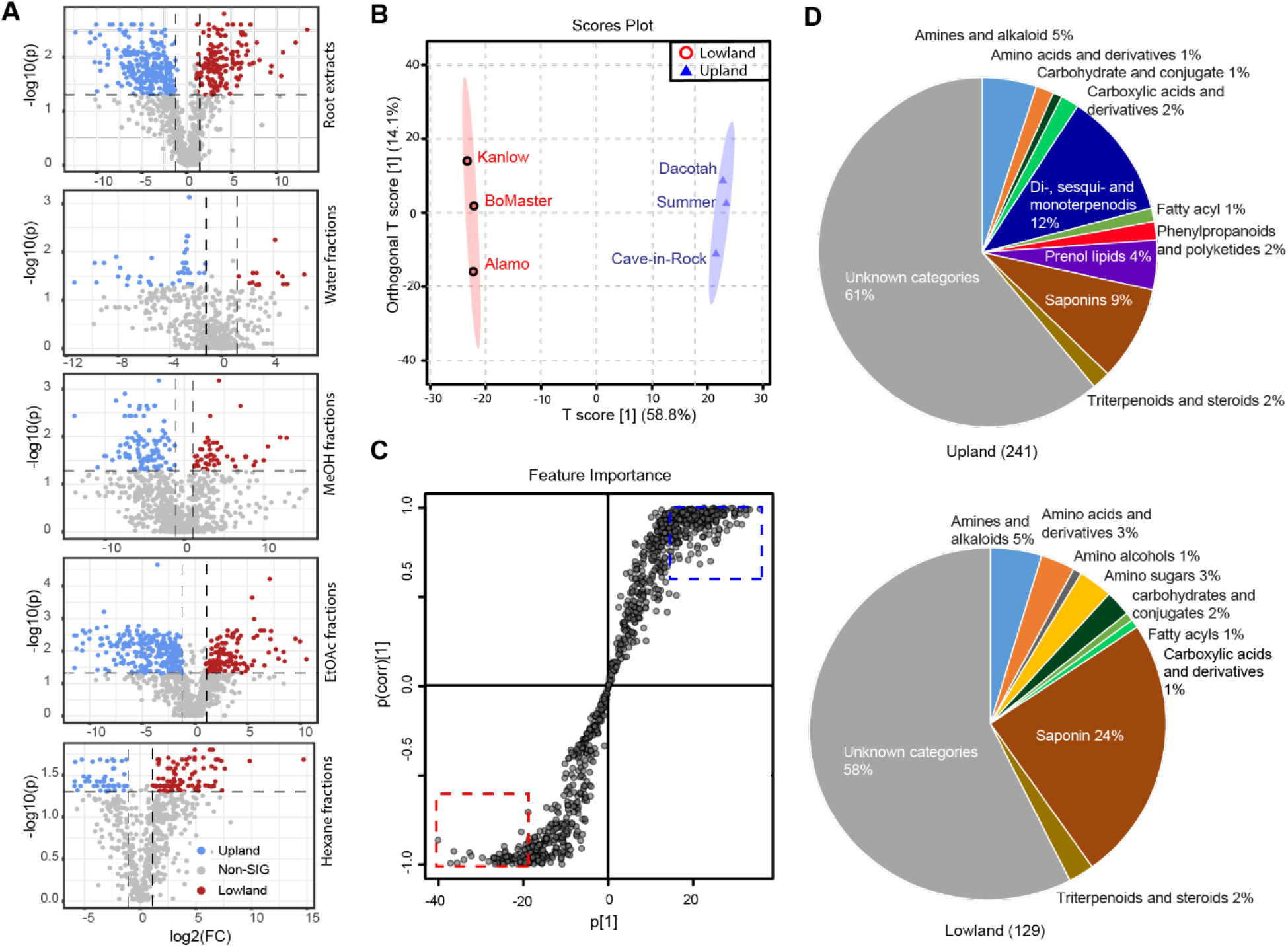
Comparative metabolomics of the upland and lowland switchgrass. (A) Volcano plots showing the results of significance analysis (cutoff threshold: FDR adjusted p ≤ 0.05; fold changes ≥ 2) that were used to identify the significant different (SIG) metabolites between the upland and lowland switchgrass ecotypes in the whole root extracts, water, methanol, EtOAc and hexane fractions, respectively. n = 3 samples each. (B) OPLS-DA scores plot of upland (blue triangles) and lowland (red circles) switchgrass whole root samples based on the normalized abundances (peaks areas) of 1777 metabolite features. (C) S-plot corresponding to the OPLS-DA model used to characterize the differentially accumulated features (DAFs) associated with upland (blue dotted rectangle) and lowland (red dotted rectangle) root samples. Cutoff values: covariance |p| ≥ 0.6 and correlation |p (corr)| ≥ 20 for whole root extracts. (D) Annotated compound classes for the identified DAFs in upland and lowland switchgrass root. Percentages for the shares within each pie chart and total numbers of the DAFs are shown in parentheses.

### Analysis of antifungal activities from switchgrass root tissues

The ecotype-specific differences in metabolite accumulation provided an opportunity to evaluate responses of epiphytic and endophytic fungi isolated from field grown switchgrass roots to extracts with unique compositions. Disc diffusion assays were set up using with mycelium inocula of 18 different fungal isolates (**Table S5**) and these were treated using filter paper discs containing upland or lowland switchgrass root extracts or solvent controls (**Fig 3A)**. The results showed that most of the tested fungi were inhibited in growth by upland and/or lowland root tissue extracts. Of note, the upland extracts inhibited more (17 out of 18) fungi than did the lowland extracts (11 out of 18, **Fig. 3 B – D and Fig S2**). Growth of only one fungus (*Myrothecium* spp.) was promoted by the extracts of both ecotypes (**Fig S2**). We were particularly interested in the fungi that differentially responded to the upland and lowland root tissue extracts; thus, we chose *Linnemannia elongata* (GLBRC 635), *Trichoderma* sp. (GLBRC 496) and *Fusarium* sp. (GLBRC 120) for detailed analysis. While extracts of both ecotypes suppressed the *L. elongata* growth, lowland ecotype cultivar extracts inhibited its growth more effectively when used at a comparable concentration (**Fig 3B**). In contrast, *Trichoderma* sp. And *Fusarium* sp. growth was strongly suppressed by the upland cultivar extracts, while lowland extracts were minimally effective (**Fig 3 C & D**).

**Figure 3.**
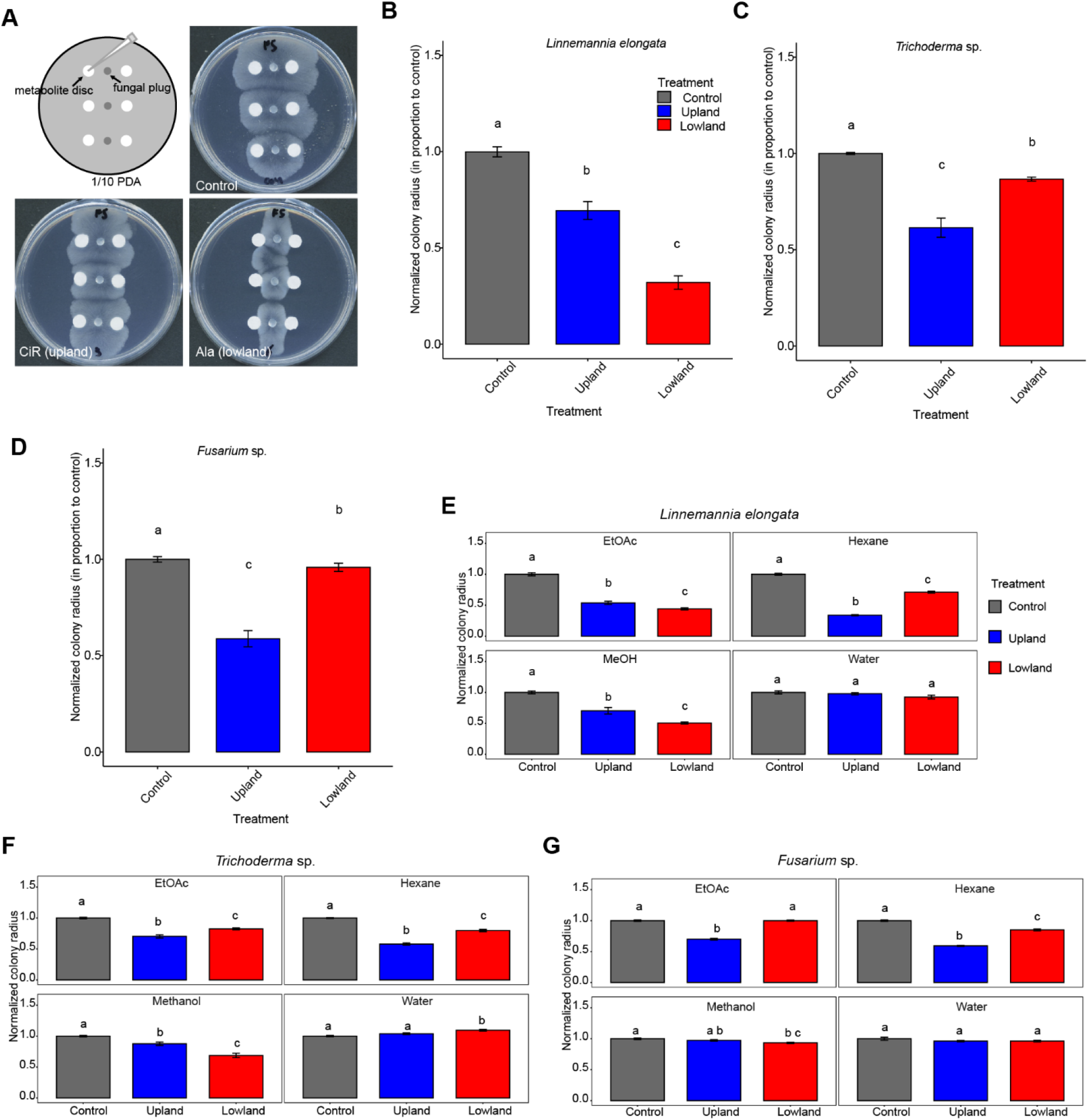
Extracts and fractions of upland and lowland switchgrass root tissues exhibit differential growth inhibition of the tested fungi. (A) Images of *L. elongata* colonies growing on the control (upper right), upland (lower left) and lowland (lower right) switchgrass root extracts in a disc diffusion assay. The fungal hyphae were inoculated along the middle vertical line at three positions; filter paper discs containing the plant extracts in ethanol were placed on both sides of each inoculation site (upper left; see **Materials and Methods** for more details). (B) – (D) Root extracts from individual upland switchgrass cultivars – Dacotah, Summer and Cave-in-Rock – and lowland cultivars – Alamo, Kanlow and BoMaster – were used to screen against *L. elongata, Trichoderma* sp. and *Fusarium* sp., respectively, at a (w/v) concentration of 50 mg/mL in 80% methanol. 80% methanol was used as the negative control. The colony radiuses from the experimental groups were normalized to the ethanol only control colony radiuses. n = 3 (cultivars) x 6 (replicates per cultivar) = 18 (total replicates). (E) – (G) Growth inhibition of different metabolite fractions from the individual upland and lowland switchgrass cultivars, used at 10 mg/mL, against *L. elongata, Trichoderma* sp. and *Fusarium* sp. 80% methanol was used as a negative control. The colony radiuses from the experimental groups were normalized to the 80% ethanol control colonies. n = 3 (cultivars) x 6 (replicates per cultivar) = 18.

We next asked whether the differences in fungal growth resulted from responses to the ecotype differentially accumulated metabolites. *Linnemannia elongata, Trichoderma* sp. and *Fusarium* sp. were first treated with fractions of upland and lowland switchgrass root tissue extracts that differed by polarity and other chemical properties (**Fig 1A**). Differential *L. elongata* growth inhibition was observed by EtOAc and hexane partitioned metabolites as well as SPE methanol fractions of the two ecotypes. The lowland EtOAc and methanol fractions were more suppressive than those of the upland cultivars, whereas the upland hexane fraction was more suppressive than the lowland one (**Fig 3E**). The *Trichoderma* sp. isolate behaved similarly to *L. elongata* except that its growth was inhibited to a greater extent by the upland EtOAc fraction (**Fig 3F**). Differential suppression by the methanol, EtOAc and hexane fractions was also observed for *Fusarium* sp.; for this isolate, the lowland methanol fraction was more suppressive than the upland methanol fraction. In contrast, the upland EtOAc and hexane fractions were more suppressive than the corresponding lowland fractions (**Fig 3G**). In contrast to the varied antifungal activities of the metabolites that partitioned into the various organic phases, none of the high-polarity metabolites enriched in the water phase inhibited the tested fungi. *Trichoderma* sp. growth was promoted slightly by the lowland water phase (**Fig 3 E – G**).

To elucidate candidate bioactive metabolites enriched in the methanol, EtOAc or hexane fractions, we performed Spearman’s rank correlation analysis to associate the MS features to antifungal performances in different root-derived fractions (**Fig 1A**). Remarkably, terpenoid specialized metabolites were highly correlated with the antifungal properties of switchgrass root extracts and fractions, revealed by the annotations of 20 MS features most highly correlated with inhibition of each of the three fungi. Five of the 20 features most inhibitory to *L. elongata* were annotated as saponins (**Fig 4**). For example, the top correlated feature was a previously unidentified steroidal saponin glycosylated at a single core position whose MS/MS spectra revealed evidence for the presence of two hexoses one of which is acetylated (**Fig S3, Table S6**). In contrast, diterpenoids were shown to be among six and eight of the most highly inhibitory metabolite features against *Trichoderma* sp. and *Fusarium* sp., respectively (**Fig 4**). Moreover, the candidate bioactive saponins and diterpenoids differed in root accumulation between the two switchgrass ecotypes; the saponins accumulated at higher levels in the lowland ecotype whereas the diterpenoids were more abundant in the upland root (**Fig 4**). These correlative results led us to test the hypothesis that the contrasting antifungal properties of switchgrass root extracts derived from these terpenoids.

**Figure 4.**
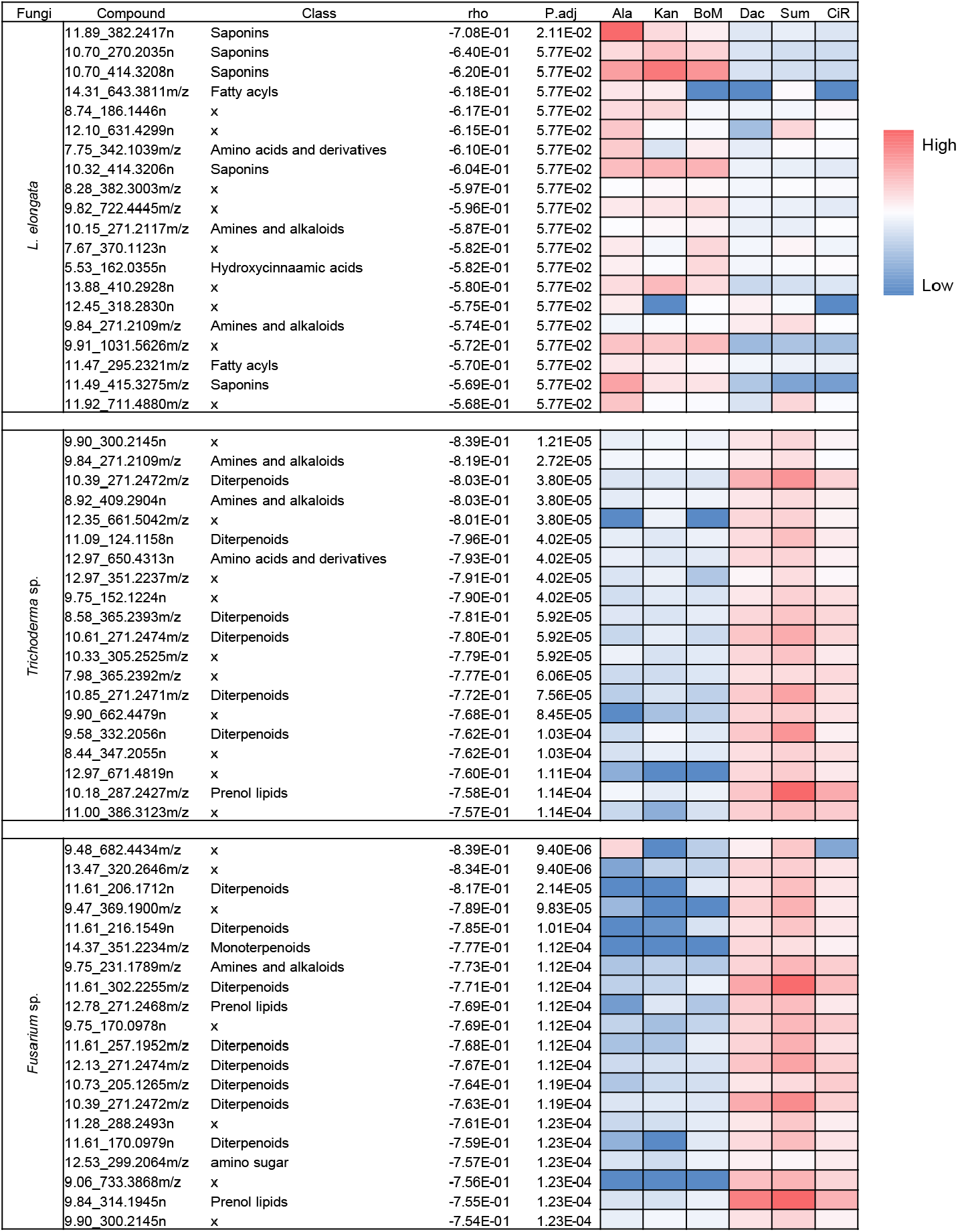
The top 20 most potent antifungal metabolites in switchgrass root tissue extracts revealed by correlation analysis. Spearman’s rank correlation coefficient was calculated to show the strength of relationship between the feature abundance and fungal growth. The coefficient, ‘rho’, has a value between 1 and -1. The features that are most negatively correlated to fungal growth are the best candidates for those contributing most to the antifungal activities of the switchgrass root extracts. The significance of the Spearman’s rank correlation coefficients was also calculated, reflected by the FDR-adjusted *p*-values. Compound classes were predicted by either CANOPUS or database searching through the Progenesis QI. ‘x’ means no compound class information associated with the feature. Features that are high in the switchgrass roots are in red, those that are low are in blue. Lowland ecotypes: Ala, Alamo; Kan, Kanlow; BoM, BoMaster. Upland ecotypes: Dac, Dacotah; Sum, Summer; CiR, Cave-in-Rock.

### Saponin and diterpenoid antifungal properties and structure – activity relationships (SARs)

To test this hypothesis, we further fractionated several structurally distinct saponins and diterpenoids from root tissues of upland or lowland switchgrass using reverse phase semi-preparative HPLC, testing their antifungal activities using the disc diffusion assay. While the resultant HPLC samples contained multiple saponins or diterpenoids, they shared the same terpene cores with differences in modifications and stereo/structural chemistry (**Table S6**). Taken together, these fractions included four classes of saponins (D415, D415-SCG, D431 and D457), which are different in their sapogenin aglycones and numbers of sugar chains, as well as three classes of diterpenoids (Di287, Di301 and Di317) that vary in their terpene backbones (**Fig 5A, Table S6**).

**Figure 5.**
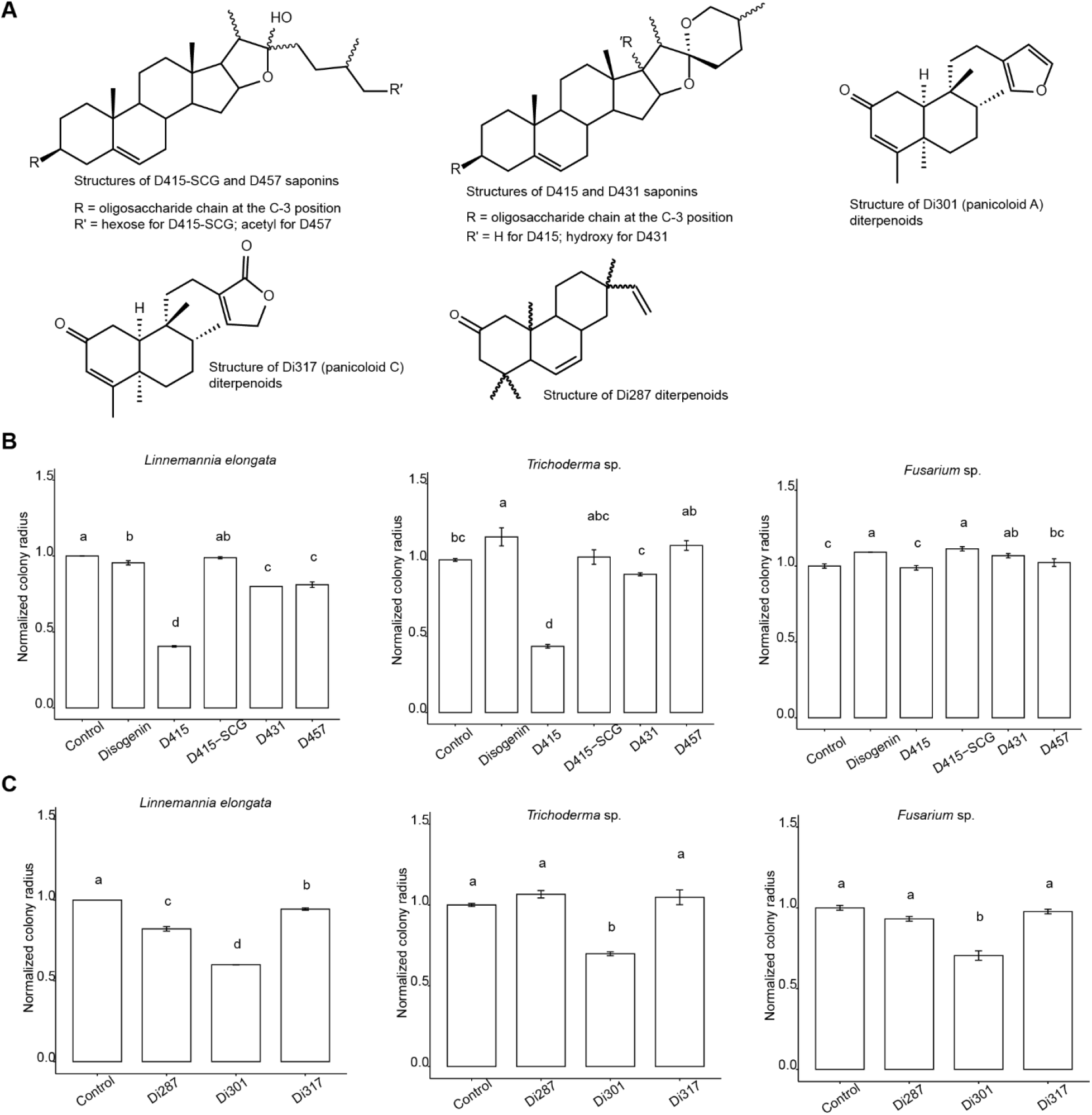
Efficacy of saponins and diterpenoids against the switchgrass rhizosphere fungal growth revealing structure-activity relationships. (A) The structures of HPLC purified switchgrass saponins and diterpenoids used in the bioassays. The structures of the saponins D415-SCG, D415, D431 and D457, and the diterpenoids, Di301 and Di317, were previously elucidated by LC-MS/MS and NMR (Li et al., 2022; Tiedge et al., 2022). NMR data supporting the structure of diterpenoid Di287 can be accessed in the supplementary files (**Fig S 4 – 12, Table S7)**. (B) Bioactivities of different saponin types were screened against *L. elongata, Trichoderma* sp. and *Fusarium* sp. in the disc diffusion assay described in Figure 3A. 50% methanol was used as the negative control. (C) Bioactivities of different diterpenoids were screened against the three tested fungi in a disc diffusion assay. Pure methanol was used as the negative control. For both the saponin 1 and diterpenoid compounds, the tested concentration was 1 mg/mL. The colony radiuses from the experimental groups were normalized to those of methanol control colonies, which were set as the scale 1 on the bar plots (n = 6).

For the saponins, we examined the activity of commercial diosgenin (a sugarless, or aglycone, sapogenin core for all the currently identified switchgrass saponins) against *L. elongata, Trichoderma* sp. and *Fusarium* sp. isolates. Consistent with the hypothesis that the sugar moieties are required for saponin anti-microbial activities (Augustin *et al*., 2011), we found no inhibitory effect with diosgenin. In contrast, we observed inhibition of *L. elongata* and *Trichoderma* sp. with the D415 saponins glycosylated at a single core position. Interestingly, the D415-SCG saponins which are glycosylated at two core positions were shown not to be inhibitory, revealing that the presence of an additional sugar moiety at the saponin backbone C-26 position resulted in loss of detectable antifungal activity (**Fig 5B**). Structural specificity for fungal growth inhibition by the saponins glycosylated at a single core position was supported by the observation that D431 and D457 (with either the C-17 hydroxylation or sidechain C-26 acetylation, respectively) showed dramatically decreased bioactivities compared to the unmodified D415 (**Fig 5B**). Diterpenoids also exhibited differences in SAR. All three fungi were inhibited by the Di301 panicoloid A and B furanoditerpenoids (**Fig 5C**) (Tiedge et al., 2022). In contrast, only *L. elongata* was also sensitive to the other two tested diterpenoids. One of these is the Di317 furanoditerpenoid panicoloid C (Tiedge et al., 2022), whose furan ring is oxidized to a ketone group (**Fig 5 C**). The other one, Di287, is a previously structurally uncharacterized and drought-inducible diterpenoid (Tiedge et al., 2022). We used a set of one- and two-dimensional NMR experiments to resolve its structure (**Materials and Methods**), revealing that this metabolite as an abietane diterpenoid without a furan ring (**Fig 5C, Fig S 4 – 12, Table S7**).

The effect of the saponins D415-SCG and D415 and the diterpenoids Di301 and Di317 on growth of *L. elongata* was examined in liquid cultures at three concentrations (0.1, 0.005 and 0.00025 mg/mL). Those results were generally consistent with the agar-based growth assay. The D415 saponin glycosylated at a single core position clearly inhibited fungal growth in a dose-dependent manner in two independent tests, while the growth inhibition by the D415-SCG saponin glycosylated at two core positions was only seen at the highest concentration in one test (**Fig S13A**). The diterpenoids Di301 and Di317 inhibited fungal growth at the highest concentrations in the first test and showed dose-dependent inhibition in the second test (**Fig S13B**). Together, these results revealed antifungal activity differences among the structurally distinct switchgrass terpenoids.

## Discussion

Switchgrass, an important constituent of the North American tallgrass prairie ecosystem, is valued as a biofuel crop for its high net energy yield and relatively low economic and environmental costs associated with irrigation, fertilizers and pesticides (Casler, Vogel and Harrison, 2015; Langholtz, Stokes and Eaton, 2016). We are interested in the possible roles of specialized metabolites in mediating documented differences in a variety of biotic and abiotic stress tolerances between the northern upland and southern lowland ecotypes of this species (Cornelius & Johnston, 1941; Nielsen, 1947; Stroup et al., 2003; Lowry et al., 2014). Here, we tested the bioactivities of upland and lowland root metabolites on 18 switchgrass rhizosphere associated fungal species, demonstrating that the steroidal saponins and diterpenoids influence ecotype-specific growth inhibition of these fungi.

Untargeted metabolomics revealed distinctive metabolomes for upland and lowland switchgrass roots (Li et al., 2022). Classification of the differentially-accumulating mass features revealed that terpenoids are among the most prominent ecotype specific metabolites. Notably, a variety of unglycosylated mono-(C10), sesqui-(C15) and diterpenoids (C20) were identified accounted for 12% of upland switchgrass DAFs while none of these metabolites were identified among the lowland ecotype DAFs (**Fig 2D, Table S 3 & 4**). In contrast, saponins – glycosylated phytosteroids or triterpenoids containing backbones of 27 - 30 carbons – were enriched among the lowland switchgrass specific metabolites. Specifically, 24% of lowland and 9% of upland ecotype DAFs were annotated as saponins (**Fig 2D, Table S 3 & 4**). Therefore, the metabolomics analysis suggested that diterpenoids and saponins are characteristic of upland and lowland switchgrass root metabolomes, respectively.

To evaluate the bioactivities of upland and lowland switchgrass root metabolites, we screened upland and lowland switchgrass root extracts for their growth impacts on 18 diverse fungal species previously isolated from switchgrass rhizosphere as epiphytes or endophytes (Beschoren da Costa *et al*., 2022). These fungi were chosen based on their prevalence in switchgrass associated environments and potential ecological roles (Beschoren da Costa *et al*., 2022). Growth of 17 of 18 fungi was inhibited by the C10, C15 and C20 terpenoid rich upland root extracts while saponin-dominated lowland root extracts suppressed growth of 11/18 isolates. Sixteen of the 18 tested fungi (16/18) differentially responded to the upland and lowland root extracts (**Fig 3 B – D and Fig S2**). Correlation analysis of these results led us to hypothesis that these fungi are more sensitive to unglycosylated terpenoids or saponins.

To pursue the hypothesis, we fractionated switchgrass root extracts based on polarity and screened representatives of three ecologically important fungal taxa, *L. elongata, Trichoderma* sp. and *Fusarium* sp., with these fractions. *Fusarium* is a prominent fungal genus in plant and soil environments (Summerell et al., 2001; Edel-Hermann et al., 2015), and the genus includes plant pathogens of economic importance (Agrios, 2005), as well as rare documented cases of beneficial effects where non-pathogenic strains of *Fusarium* compete with pathogenic *Fusarium* in diseases such as *Fusarium* wilt (Steinberg *et al*., 2019). *Trichoderma* isolates were documented to be beneficial, pathogenic or commensal. As one of the most prevalent fungi in soil and plant environment (Harman and Kubicek, 2002), they were also found to be plant symbionts that promote plant growth and defense (Naseby et al., 2000; Harman et al., 2004; Hajieghrari et al., 2008; Guzmán-Guzmán et al., 2019). However, a few species were also found to be pathogenic to plants (Munkvold and White, 2016; Pfordt *et al*., 2020). *Linnemannia*, formerly classified as *Mortierella*, is closely associated with switchgrass roots (Vandepol *et al*., 2020; Beschoren da Costa *et al*., 2022). Isolates of the species promoted growth of *Arabidopsis* and economically important crops such as *Citrullus lanatus, Zea mays, Solanum lycopersicum, Cucurbita* and *Calibrachoa* through inducing IAA production and facilitating P and Fe acquisition (Li *et al*., 2018; Becker and Cubeta, 2020; Zhang *et al*., 2020; Ozimek and Hanaka, 2021; Vandepol *et al*., 2022).

We tested the bioactivities of upland and lowland switchgrass root fractions with different metabolite compositions against these three focal fungal species. Overall, we observed differential responses of these species to the saponin enriched methanol fraction and unglycosylated terpene enriched EtOAc/hexane fractions. For example, growth of *L. elongata* was more sensitive to the lowland methanol fraction whereas the upland EtOAc/hexane fractions inhibited *Trichoderma* sp. and *Fusarium* sp. (**Fig 3 E – G**). Furthermore, correlation analysis (**Fig 4)** predicted that growth of the three focal species was most highly influenced by saponins and diterpenoids.

We tested the bioactivities of these terpenoids by using a set of structurally distinct steroidal saponins and diterpenoids purified from upland and lowland switchgrass roots (**Fig 5**). This SAR analysis revealed that the saponin antifungal activity was primarily associated with the number of saponin sugar chains: our results revealed that the saponins glycosylated at a single core position were inhibitory whereas those glycosylated at two core positions were not (**Fig 5B**). The C-3 sugar substitution seems to be necessary for the antifungal activities as the aglycone sapogenin, diosgenin, only minimally inhibited the growth of *L. elongata*, and even slightly promoted growth of *Trichoderma* sp. and *Fusarium* sp. (**Fig 5B**). In addition, the C-17 hydroxylation and C-26 acetylation dramatically reduced the antifungal activities of saponins with a single sugar chain (**Fig 5B**). A similar observation was reported for oat avenasides, where the 26-desgluco-avenacosides inhibited *Sordaria macrospora* growth, while the intact avenacosides showed no effect at the same or twofold higher concentration (Nisius, 1988). These results are consistent with the hypothesis that fungal growth inhibition by saponins acts by their hemolytic activity – the same molecular mechanism causing lysis of mammalian erythrocytes (Bangham and Horne, 1962; Seeman, Cheng and Iles, 1973; Baumann *et al*., 2000; Augustin *et al*., 2011). Such a mechanism requires both hydrophilic sugar chains and non-polar end of sapogenin core for formation of aqueous pores leading to membrane perturbation (Keukens et al., 1995; Armah et al., 1999; Lin & Wang, 2010).

We purified and elucidated the structure for a novel abietane type diterpenoid – Di287 – from switchgrass roots. In contrast to the previously identified Panicoloid diterpenoids (Di301 and Di317, Tiedge et al., 2022), the NMR evidence indicated lack of a furan moiety in the diterpene scaffold of Di287 (**Fig S 4 – 12**). We then explored SAR of three different types of switchgrass diterpenes – Di301, Di317 and Di287 – with and without a furan moiety. In contrast to the strong structural specificity of saponins, all three types of diterpenoids had antifungal activities against *L. elongata* (**Fig 5C**). Di301, containing a furan moiety in its side chain, exhibited the strongest growth inhibition. It also inhibited growth of *Trichoderma* sp. and *Fusarium* sp., while Di287 – lacking the furan moiety – and the oxidized furan ring containing Di317 did not (**Fig 5C**). These observations indicate that a furan ring increases the switchgrass diterpenoid antifungal activities, while oxidation of this moiety results in lower potency. Regardless of the structural characteristics, significantly increased accumulations of all three diterpenoid types were observed in the roots of drought stressed switchgrass (Tiedge et al., 2022), suggesting potential functions of these structurally diverse terpenoids beyond their antifungal activities.

In conclusion, we demonstrated differential inhibition of ecologically important fungi isolated from the rhizosphere of the perennial bioenergy crop by saponins and diterpenoids – the metabolites preferentially accumulated in the upland and lowland switchgrass roots, respectively. Results of this study add to published work demonstrating that the antimicrobial properties of plant specialized metabolites influence microbiome assembly in crops as diverse as *Panax notoginseng* (Wei et al., 2022), *Camellia oleifera* (Zhang et al., 2022), *Lilium Oriental* (Liu *et al*., 2011), and *Zea mays* (Murphy *et al*., 2021). Together with these previous studies, our results suggest that ecotypic differences in switchgrass terpene specialized metabolites might differentially influence soil fungal microbiome activities. This study sets the stage for further investigation of the *in vivo* roles of these specialized metabolites in assembling switchgrass associated microbial communities and developing breeding strategies to facilitate biomass production and sustainable cropping system for switchgrass.

## Material and Methods

### Plant materials

Seeds for the upland (Dacotah, Summer and Cave-in-Rock) and lowland (Alamo, Kanlow, and BoMaster) switchgrass were purchased from Native Connections (http://nativeconnections.net, Three Rivers, MI). The seeds were sowed into a 1:1 mixture of sand and vermiculite and grown for three months at 27 °C with 16 h light (cool white fluorescent light bulbs, 500 μE m^-2^s^-1^) per day in a growth chamber. The relative humidity in the chamber was set to 53% without dehumidification.

### Metabolite extraction and fractionation

All chemicals were obtained from Sigma Aldrich (St. Louis, MO) unless otherwise specified. Switchgrass plants were pulled out of pots and cut ∼1 cm above the rhizome to collect the root tissues. Attached sand and vermiculite were carefully washed from the root surface using deionized water and blotted dry by paper towels. 100 – 200 g fresh switchgrass root tissues were chilled in liquid nitrogen and powdered using mortar and pestle. Total metabolites were obtained by immediately adding 80% methanol to the powders at a tissue/solvent ratio 1:10 (w/v) and extracting (without stirring) at 4° C overnight. The solvent was removed and the extraction was repeated with fresh 80% methanol at 4° C overnight. The combined solvent was centrifuged at 4000 g for 15 min to remove the debris, pooled, concentrated using a rotary evaporator and then completely dried down in a SpeedVac concentrator (Thermo Scientific, Waltham, MA). The dried extracts were stored in a -80° C freezer.

Sequential liquid-liquid phase partitioning and solid phase extraction (SPE) for the whole root extracts was carried out according to a published protocol (Ebada *et al*., 2008). Briefly, the dry extracts were dissolved in 100 mL water and phase partitioned against hexane at a 1:1 ratio in a separatory funnel. The water phase was next phase partitioned against EtOAc. Each of these liquid-liquid phase partitioning steps was done only once. After that, the resultant water phase was loaded to a 35 cc C18 SPE cartridge (Waters, Milford, MA). The cartridge was first washed using 3 × 20 mL of each 0%, 10% and 20% methanol (in water). The flowthrough and wash were collected, pooled and considered as the final ‘water phase’ in this study. The cartridge was then eluted again using 3 × 20 mL of each 50%, 70%, 80% and 90% methanol (in water). These washes were collected, pooled and called the ‘methanol phase’. All fractions were evaporated to dryness under vacuum using a SpeedVac and stored in the -80 freezer before bioassay.

### Compound purification

The methanol fraction and EtOAc/hexane fraction were used for steroidal saponin and diterpenoid isolation via reverse phase HPLC, respectively. Steroidal saponins were purified from the methanol fraction of lowland switchgrass, Kanlow, root extracts, following the procedures previously described (Li *et al*., 2022). Diterpenoids were purified from the EtOAc and hexane fractions of upland switchgrass, Cave-in-Rock, root extracts. Diterpene purification was performed using a Waters 2795 pump/autosampler connected with LKB Superrac 2211 fraction collectorand a Waters Symmetry C18 HPLC column (100 Å, 5µm, 4.6 mm x 150 mm). The water mobile phase was adjusted to PH 2.8 (Solvent A) and acetonitrile was used as the organic mobile phase (Solvent B). A 20-min linear gradient elution was used to purify all three diterpenoids, Di287, Di301 and Di317, starting from 1% B at 0 min, 40% B at 1.01 min and held at 40% between 1.01 and 3 min, linear increased to 90% at 15 min, 99% B at 15.01 min and held at 99% B between 15.01 and 18 min. The solvent flow rate and column temperature were set at 1.5 mL/min and 40°C, respectively. Followed by a sample injection (100 µL), the 20-min LC gradient starts during which liquid droplets were collected, by using a fraction collector, every 10 secs as a single fraction. In the end, 120 fractions were obtained. The HPLC fractions containing the same targeted compounds, viewed by LC-MS, were pooled and dried down under vacuum in a SpeedVac.

### LC-MS metabolite analysis

For each LC-MS sample preparation, 500 mg fresh switchgrass root tissues were powered with liquid nitrogen using on a Mini G homogenizer (SPEX SamplePrep, Metuchen, NJ) and extracted in 5 mL 80% methanol containing 1 μM telmisartan internal standard and incubated for 16 h at 4°C (without stirring). Samples were centrifuged for 20 min at 4000 g and the supernatants transferred into 2-mL HPLC vials and stored in a -20°C freezer prior to LC-MS analysis. The switchgrass metabolite separation and MS analysis were done using a reversed-phase UPLC BEH C18 (2.1 mm x 150 mm, 1.7 μm column, Waters) connected with an Electrospray Ionization – Quadrupole Time-of-Flight MS (ESI-QToF-MS, Waters). Both positive and negative mass spectral information were acquired under data-independent acquisition (DIA, MS^E^) and data-dependent acquisition (DDA, MS/MS) modes. The later method was specifically used for feature annotation purpose. All LC gradient and MS condition experimental parameters were as previously reported (Li *et al*., 2022)

### Untargeted metabolomics data processing

The raw MS data pre-treatments, including peak picking adduct ion grouping, retention time (RT) alignment and lock mass correction, were performed in the Progenesis QI software (v3.0, Waters). By doing so, the metabolite features were identified, and each was defined by its RT and mass-to-charge (*m/z*) value. The intensities (abundances) of features were calculated from the peak area information, and then filtered to choose the ones with abundances ≥ 500 for the downstream statistical analysis. The parameters used with Progenesis processing were same as described in Li et al. (2022). The untargeted metabolomics feature annotation was achieved in two separate ways. First, online databases, including MassBank, MetaboLights, Plant Metabolic Network and PlantCyc were used to in silico search for the matching compounds deposited in the databases with the features identified from switchgrass, based on 10 *ppm* precursor tolerance, 95% isotope similarity and 10 *ppm* theoretical fragmentation pattern matching with fragment tolerance. Second, the MS/MS data acquired by DDA were used for chemical class predications for the untargeted metabolomics features using the CANOPUS (class assignment and ontology prediction using mass spectrometry) machine learning function (Dührkop *et al*., 2021) built in the SIRIUS 4, which is a java-based software framework for the analysis of LC-MS/MS data of metabolites (https://bio.informatik.uni-jena.de/sirius/).

### NMR Spectroscopy

For NMR analysis of the Di287 diterpenoid, the HPLC purified sample was dried under vacuum and dissolved in deuterated chloroform (CDCl3). NMR spectra were acquired using a Bruker Avance NEO 600 MHz spectrometer (Bruker Biospin, Germany) operating at 600.13 MHz for proton experiments and equipped with a 5 mm nitrogen cryogenic HCN Prodigy probe and a SampleCase autosampler with sample cooling capability to 6 °C. For ^1^H NMR spectra, solvent suppression with shaped pulse program (wetdc) was used with a scan number (ns) of 64 at temperature 298K with pulse width 8 µs and power 10.3 W. Acquisition time for each scan was 0.681 s with a 10 s delay time for a spectral width (sw) of 19.8 ppm. The magnetic field was locked to the solvent, CDCl3 and ^1^H spectra were calibrated using the residual solvent peaks. For the subsequent 1D and 2D NMR experiments, the spectra were collected using default Bruker pulse program jresqf for HOMO2DJ (ns= 16, sw= 19.8), zgpg30 for ^13^C (ns = 10,000, sw= 236), cosygppqf for COSY (ns= 8, sw= 13 for F1 & F2), hsqcedetgpsisp2.3 for HSQC (ns= 16, sw= 13 for F2 and 220 for F1), hmbcetgpl3nd for HMBC (ns= 32, sw=13 for F2 and 220 for F1), h2bcetgpl3 for H2BC (ns= 32, sw= 15.1 for F2 and 180 for F1) and mlevphpp for TOCSY (ns=64, sw= 13 for F1 & F2) with pulse width 8 µs and power 10.3 W for ^1^H and pulse width 11.5 µs and power 152W for ^13^C. All NMR spectra were acquired with Topspin 3.5.6. (Bruker) and analyzed using MestReNova 14.2.0 software (Mestrelab Research, Spain). Chemical shifts were normalized to the chloroform NMR solvent signals of ^1^H 7.26 *ppm* and ^13^C 77 *ppm*.

### Fungal bioassay

There were in total 18 fungal isolates (**Table S5**) used in this study which were obtained from switchgrass roots as previously described (Beschoren da Costa et al., 2022). The initial fungal growth assays were performed on all 18 isolates with bulk root extracts using a disc diffusion method. For root extracts from each switchgrass cultivar, six replicates were conducted on a 5 cm radius Petri dish containing 0.1× PDA, comprised of potato dextrose broth mix (BD Diagnostics, Sparks, MD) at 2.4g/L concentration, agar at 1.5% concentration amended with streptomycin and kanamycin at 50 µg/mL concentration. To commence the assay, 8 µL root extracts pre-dissolved in 70% ethanol at 50 mg/mL concentration was applied to each sterilized 0.6 cm radius round filter paper discs (Whatman 4, Cytiva, Maidstone, UK), and the solvent was allowed to evaporate in a biosafety hood for 1 hr before placing on the growth media. Fungi were grown on PDA, and 2 mm radius plugs were generated with a flame sterilized metal borer. These were used to inoculate the fungal growth assay by placing the plugs 1 cm away from the paper discs on each side. Fungal species-switchgrass cultivar growth assays were compared to a control that had paper discs with 70% ethanol followed by evaporation in a biosafety hood for 1 hr. The Petri dishes were incubated under room temperature (∼23°C) and the colony horizontal growth from the center of the fungal plug to the rim of the mycelia was measured every other day.

From the initial 18 isolate screening, three isolates, *Trichoderma* sp. (GLBRC 497), *Fusarium* sp. (GLBRC 120) and *Linnemannia elongata* (GLBRC 635), with ecological importance were then chosen to proceed with the follow-up assays. The fungal growth assays on metabolite fractions from switchgrass root extracts were performed according to the above method except the paper discs were applied with metabolites dissolved in distilled water, 80% Methanol, EtOAc or hexane. The metabolite concentration (w/v) for each fraction was 10 mg/mL. The fungal growth was compared with a control grown with evaporated paper discs applied with the same solvent as the relevant fraction.

Liquid culture assays were set-up in pre-weighted 2 mL microcentrifuge tubes with 1.8 mL 0.1X PDB each containing purified Di301, Di317, D415, or D415-SCG at concentrations of 5, 0.25, and 0.0125 mg/mL, and 20 μl of *L. elongata* hyphae lysate in six replicates. Extra no metabolite positive and 0.1X PDB only negative controls were prepared with six replications to compare with the treatment, respectively. The lysate of *L. elongata* hyphae was prepared by putting approximately 100mg fresh PDB cultured *L. elongata* hyphae in 1.8 ml 0.1X PDB along with three 2 mm radius sterile metal beads, and vortaxing at maximum speed for 3 min. The liquid culture assay was incubated on an orbital shaker at 100 RPM and 25°C for 96 h before measuring the hyphae fresh weight. The hyphae fresh weights were measured by first centrifuging the 2 mL tubes containing the liquid fungal culture at 17, 000 g for 10 mins at the room temperature to pellet down the hyphae. The supernatant was removed using a P1000 pipette. The resultant tubes with fungal hyphae were weighted on a XP26 high performance analytical balance (Mettler-Toledo, Columbus, OH). The hyphae fresh weight was obtained by subtracting the weight of the empty 2 mL tube from the weight of the tube with the hyphae.

Measured fungal growth was normalized against the control to make all the metabolite treated fungal growth relative to the growth of the control. All measurements from the disc diffusion and liquid culture assays were statistically analyzed and visualized with one-way ANOVA followed by Tukey HSD in R with packages “agricolae” and “ggplot2.”

### Statistical analysis for the untargeted metabolomics analysis

Hierarchical Clustering Analysis (HCA), fold change analysis (volcano plot), Principal Component Analysis (PCA) and Orthogonal Projections to Latent Structures Discriminant Analysis (OPLS-DA) were done on the MetaboAnalyst 5.0 online tool platform. The *p*-values from all the significance analysis were adjusted by false discovery rate (FDR). An adjusted *p-* value ≤ 0.05 was considered statistically significant. The Spearman’s correlation test was conducted in R and *p*-values were adjusted with Benjamini and Hochberg false discovery rate correction method.

## Supplemental Data

The following supplemental materials are available.

**Supplemental Figure S1**. Differences in the metabolite profiles of upland and lowland switchgrass root fractions.

**Supplemental Figure S2**. Effect of switchgrass root extracts on disc diffusion assay growth of 15 switchgrass rhizosphere fungal isolates.

**Supplemental Figure S3**. MS/MS spectra annotation for the saponin, SS782, obtained by positive LC-MS analysis, DDA mode.

**Supplemental Figure S4**. Structure (A) and key COSY (red, bold) and HMBC/H2BC (blue arrow) correlations (B) for the abietane diterpenoid – Di287 – purified from the switchgrass root extracts.

**Supplemental Figure S5**. ^1^H NMR spectrum for the diterpenoid Di287.

**Supplemental Figure S6**. gCOSY ^1^H – ^1^H NMR spectrum for the diterpenoid Di287.

**Supplemental Figure S7**. gHSQCAD ^1^H – ^13^C NMR spectrum for the diterpenoid Di287.

**Supplemental Figure S8**. gHMBCAD ^1^H – ^13^C NMR spectrum for the diterpenoid Di287.

**Supplemental Figure S9**. gH2BCAD ^1^H – ^13^C NMR spectrum for the diterpenoid Di287.

**Supplemental Figure S10**. gTOCSY ^1^H – ^1^H NMR spectrum for the diterpenoid Di287.

**Supplemental Figure S11**. HOMO2DJ NMR spectrum for the diterpenoid Di287.

**Supplemental Figure S12**. ^13^C NMR spectrum for the diterpenoid Di287.

**Supplemental Figure S13**. The saponins and diterpenoids inhibited *Linnemannia elongata* growth in liquid-based bioassays.

**Supplemental Table S1**. The untargeted metabolomics dataset.

**Supplemental Table S2**. Annotations for the metabolite features based on LC-MS/MS spectra.

**Supplemental Table S3**. The upland switchgrass ecotype differentially accumulated features (DAFs) enriched in the upland root extracts and fractions identified by OPLS-DA analysis.

**Supplemental Table S4**. The lowland switchgrass ecotype differentially accumulated features (DAFs) enriched in the lowland root extracts and fractions identified by the OPLS-DA analysis.

**Supplemental Table S5**. Names and Genbank accession numbers for the 18 switchgrass fungal isolates.

**Supplemental Table S6**. Metabolite annotation and documentation for the UPLC-QTOF-MS data of switchgrass steroidal saponins and diterpenoids.

**Supplemental Table S7**. NMR chemical shift assignments of the diterpenoid Di287.

## Author Contributions

X.L.: conceived and performed research and data analysis, created figures and tables, and wrote the manuscript. M-Y.C: conceived and performed research and data analysis, created figures and tables, and wrote the manuscript G.M.B. conceived of experimental approaches and edited the manuscript. R.L.L.: conceived of experimental approaches and wrote and edited the manuscript.

## Acknowledgments

We thank Anthony Schilmiller (MSU Mass Spectrometry and Metabolomics Core) for LC–MS analysis-related technical support, and A. Daniel Jones for help with NMR analysis. We appreciate the assistance from Sophie Gabrysiak and ShuFan Lin on setting up the fungal growth assays. This material is based upon work supported by the Great Lakes Bioenergy Research Center, U.S. Department of Energy, Office of Science, Office of Biological and Environmental Research, under award No. DE-SC0018409.

