## Supplementary Figures S1-S13 for "Identification of anti-fungal bioactive terpenoids from the bioenergy crop switchgrass (*Panicum virgatum*)"

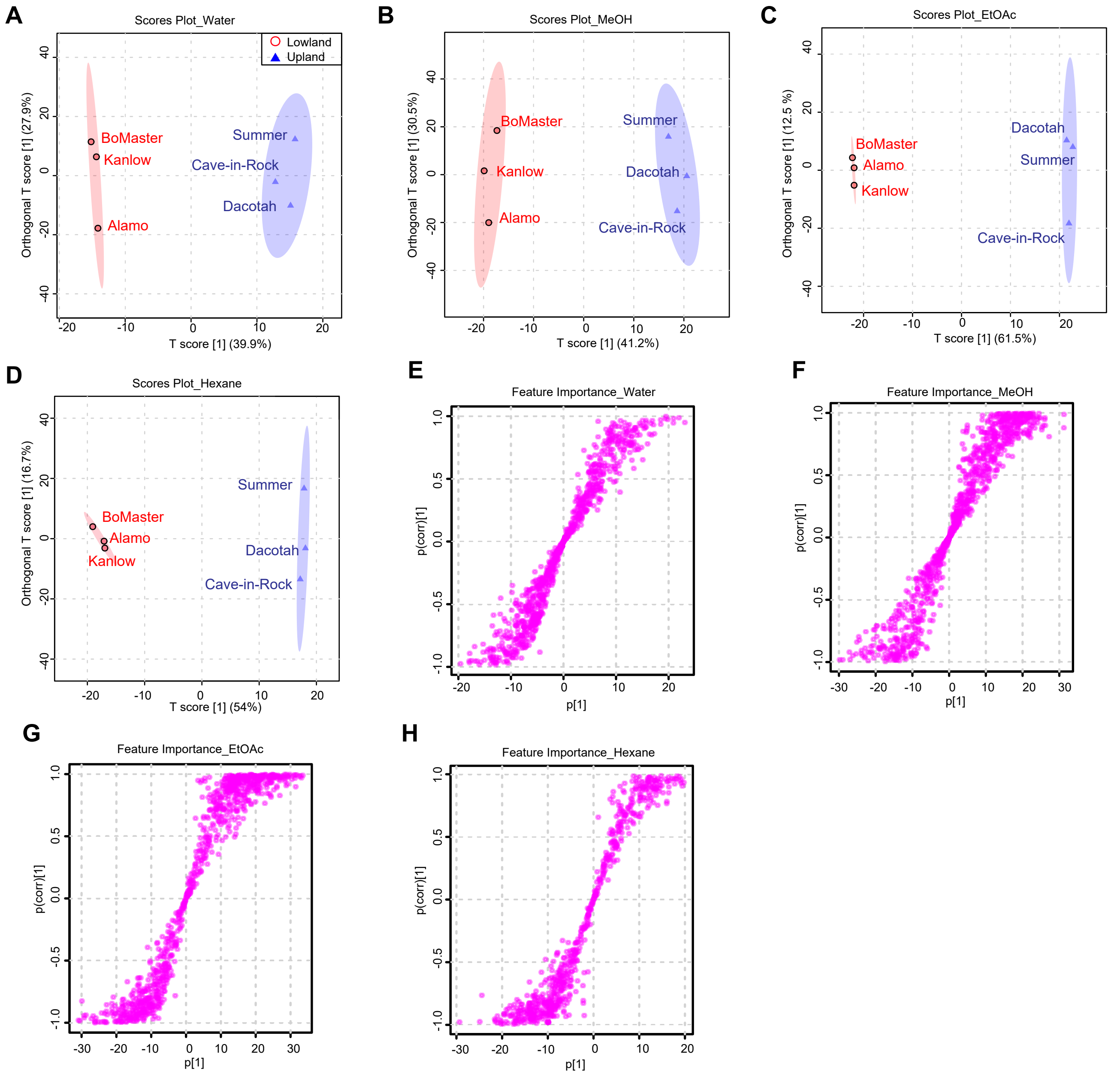


**Figure S1. Differences in the metabolite profiles of upland and lowland switchgrass root fractions.** (A) – (D) OPLS-DA score plots of upland (blue triangles) and lowland (red circles) switchgrass cultivar root fractions (water, MeOH, EtOAc and hexane) on the basis of normalized positive-mode LC-MS peak areas of 1777 metabolite features. (E) – (H) S-plots corresponding to the OPLS-DA models used to characterize the DAFs enriched in the upland and lowland root fractions. Cutoff values for the differentially accumulated features (DAFs): covariance |p| ≥ 0.6 and correlation |p (corr)| ≥ 20 for methanol and EtOA; covariance |p| ≥ 0.6 and correlation |p (corr)| ≥ 10 for water and hexane.

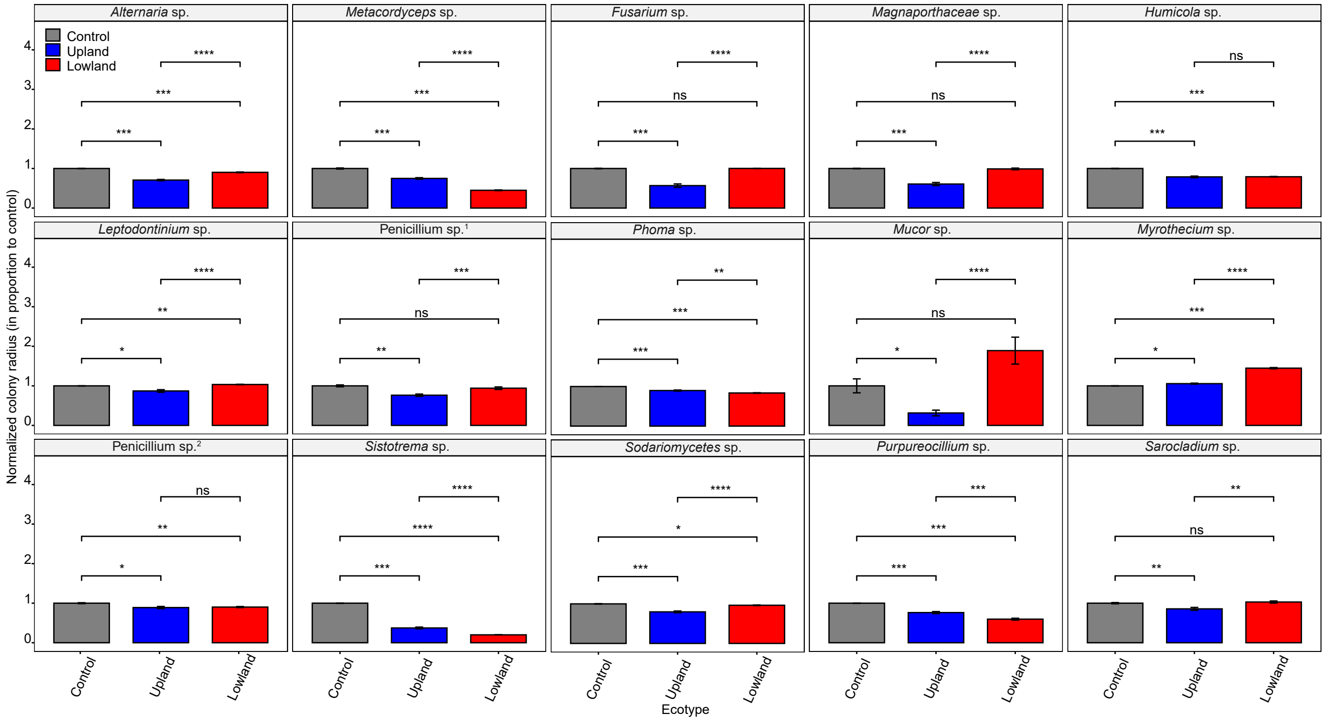


**Figure S2. Effect of switchgrass root extracts on disc diffusion assay growth of 15 switchgrass rhizosphere fungal isolates.** The upland switchgrass cultivars (blue) are Dacotah, Summer and Cave-in-Rock; the lowland switchgrass cultivars (red) are Alamo, Kanlow and BoMaster. The names of fungal isolates are shown on the top of each panel. The root extract concentrations were 50 mg/mL. 80% methanol was used as a negative control. The colony radiuses from the experimental groups were normalized to the control colony radiuses. n = 3 (cultivars) x 6 (replicates per cultivar) = 18 (total replicates). *Penicillium* sp.[1], GLBRC_165; *Penicillium* sp.[2], GLBRC_242 (see **Table S5** for more details).

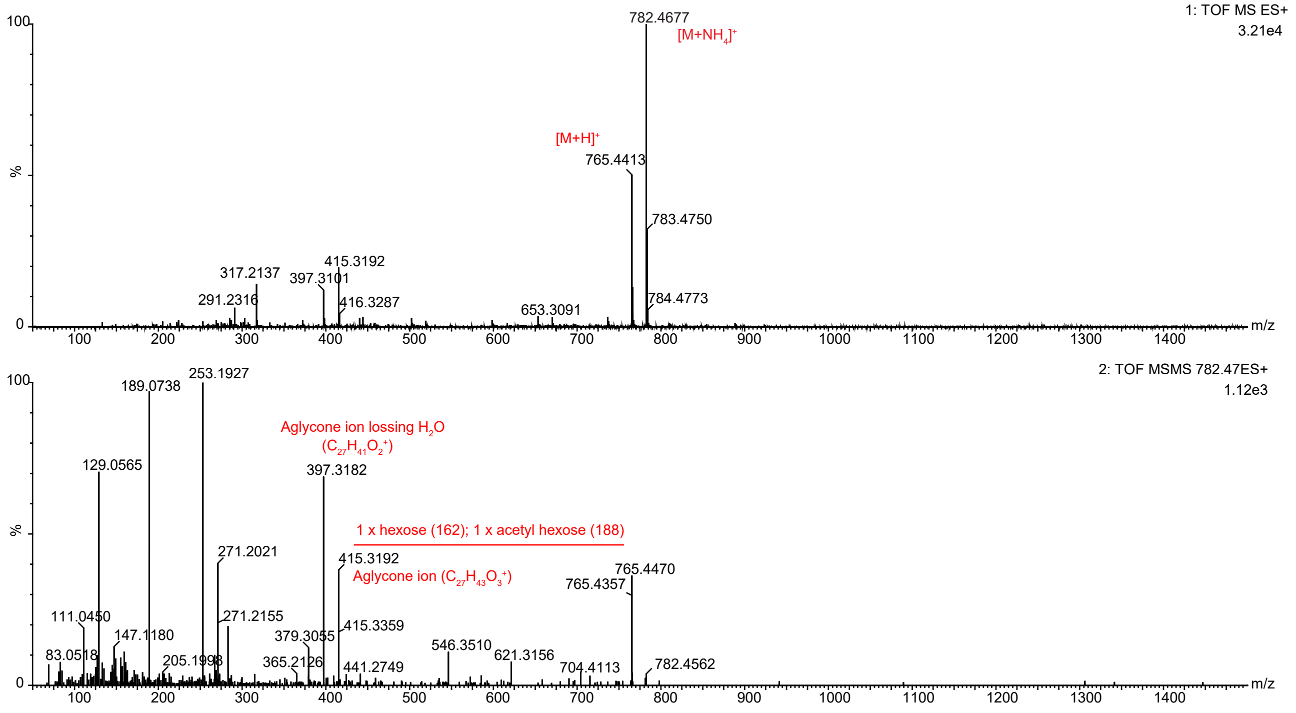


**Figure S3. MS/MS spectra annotation for the saponin, SS782, obtained by positive LC-MS analysis, DDA mode.** This saponin belongs to the monoglycosylated saponin class, D415, with a formular $C_{41}H_{64}O_{13}$ (**Table S5**). It was identified by the Progenesis QI as the feature, '11.89_382.2417n' (**Table S1, S4**) in the positive mode LC-MS analysis. Top trace: survey scan; bottom trace: MS/MS. The molecular ions, sapogenin aglycone fragment ions and neutral mass loss are indicated. 'M' standards for molecular ion.

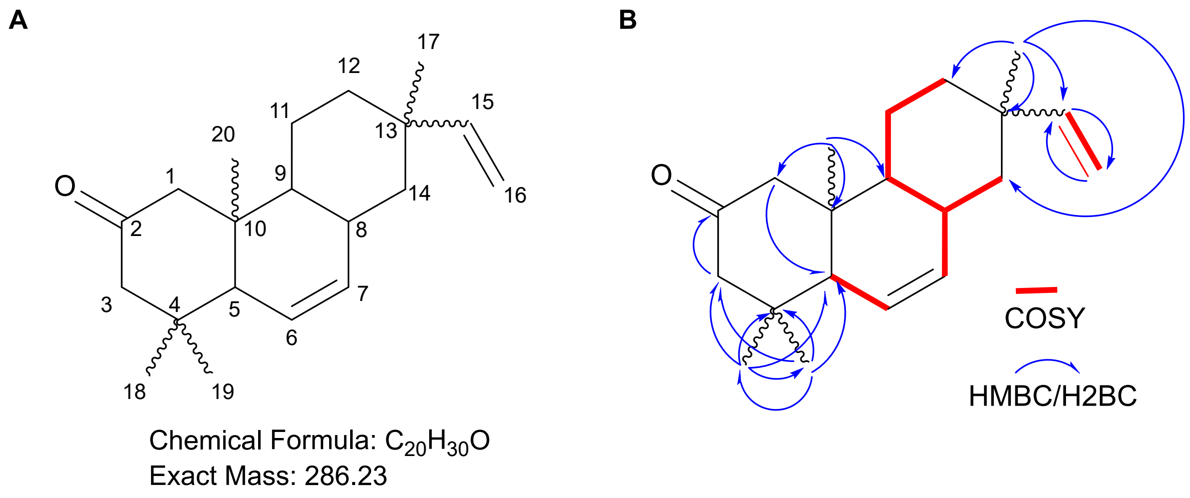


**Figure S4. Structure (A) and key COSY (red, bold) and HMBC/H2BC (blue arrow) correlations (B) for the abietane diterpenoid – Di287 – purified from the switchgrass root extracts.**

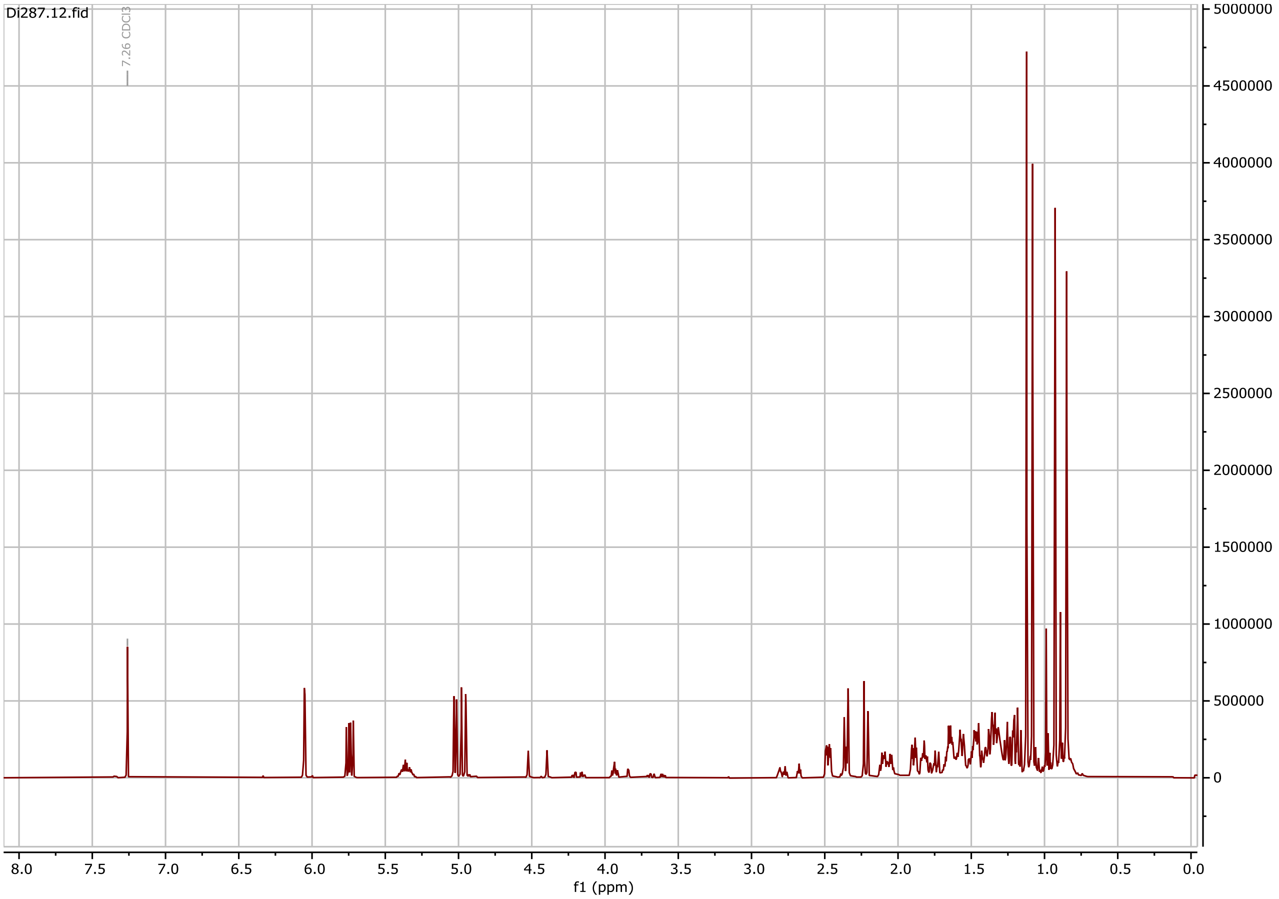


**Figure S5.** **$^{1}$H NMR spectrum for the diterpenoid Di287.**

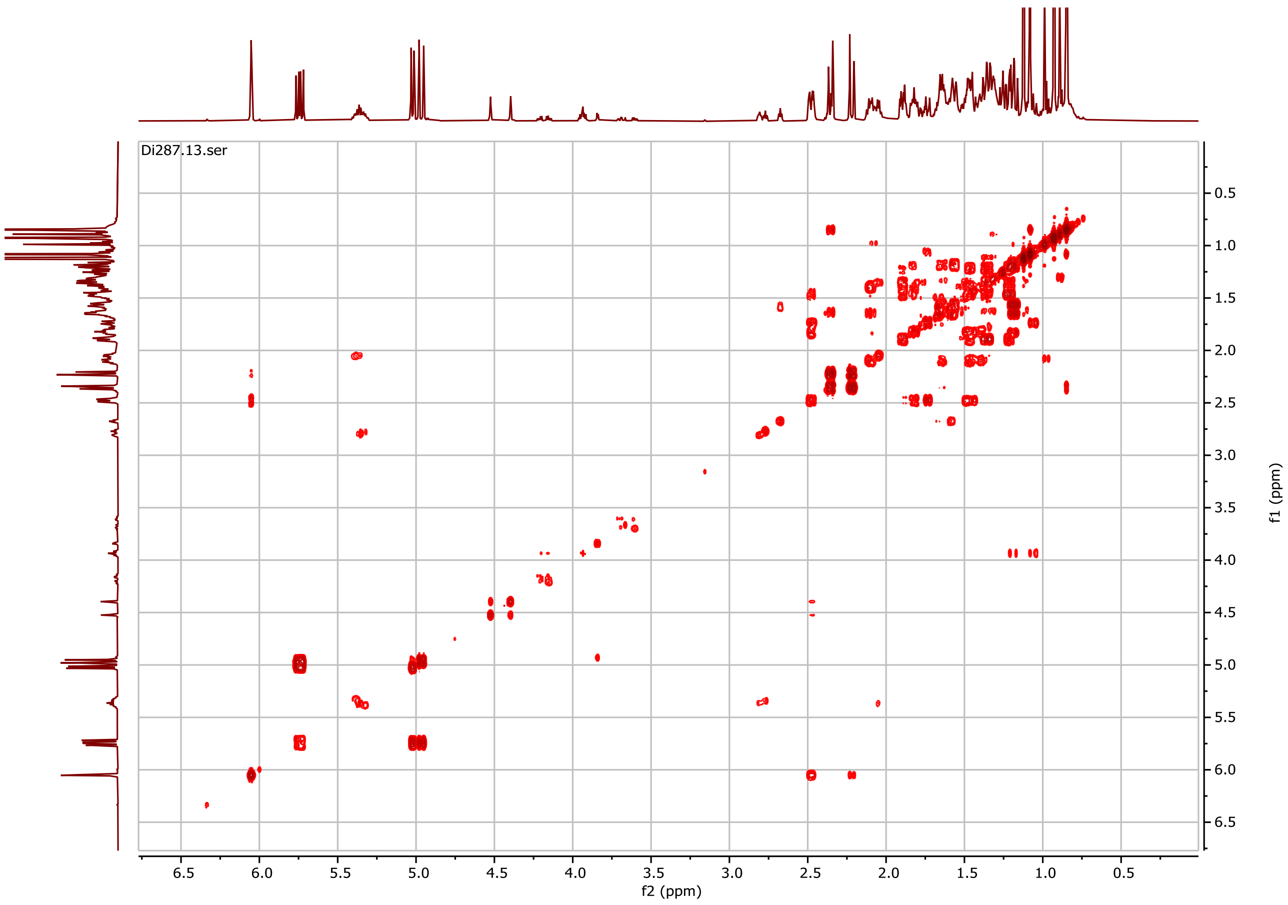


**Figure S6. gCOSY $^{1}H - ^{1}H$ NMR spectrum for the diterpenoid Di287.**

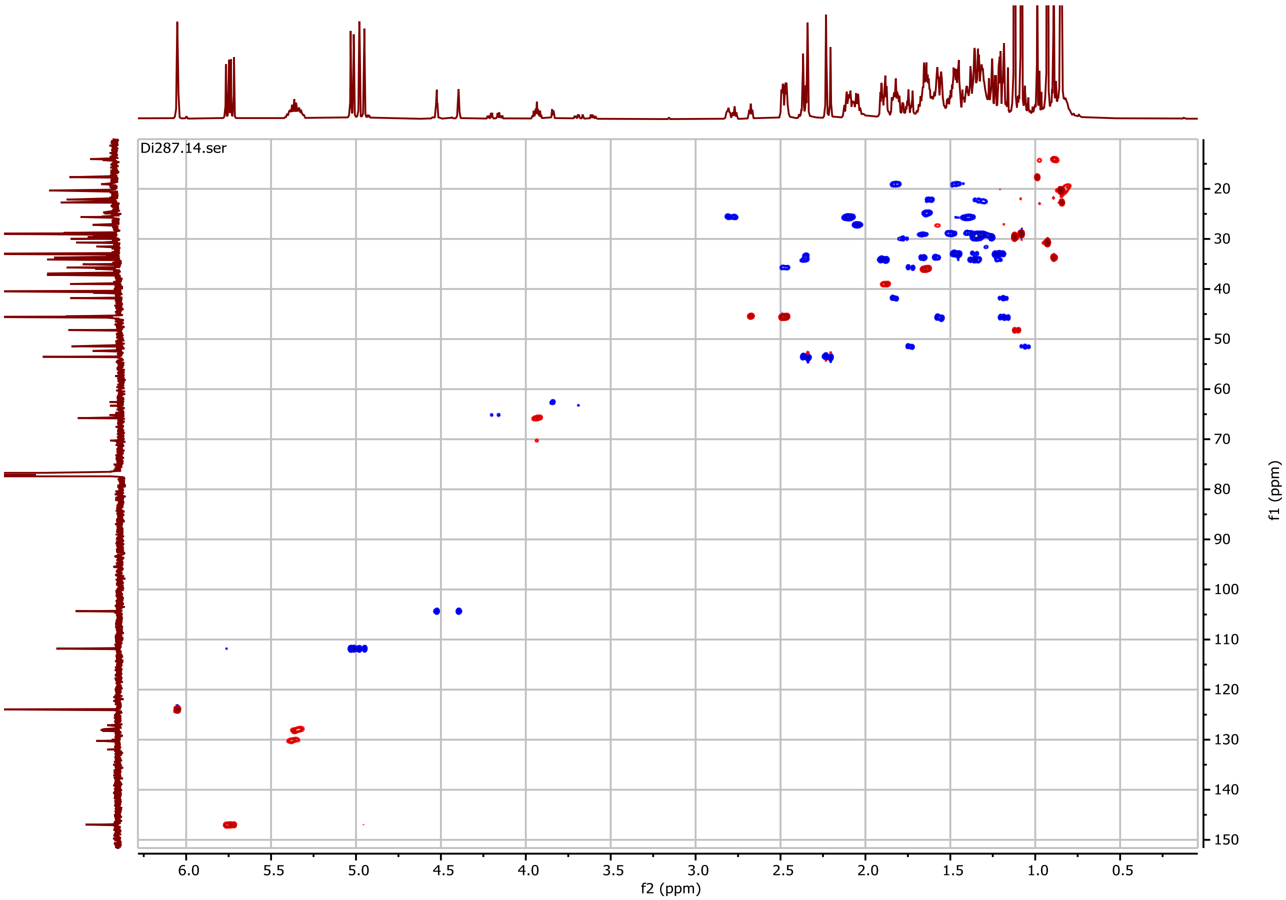


**Figure S7. gHSQCAD $^{1}H$ – $^{13}C$ NMR spectrum for the diterpenoid Di287.**

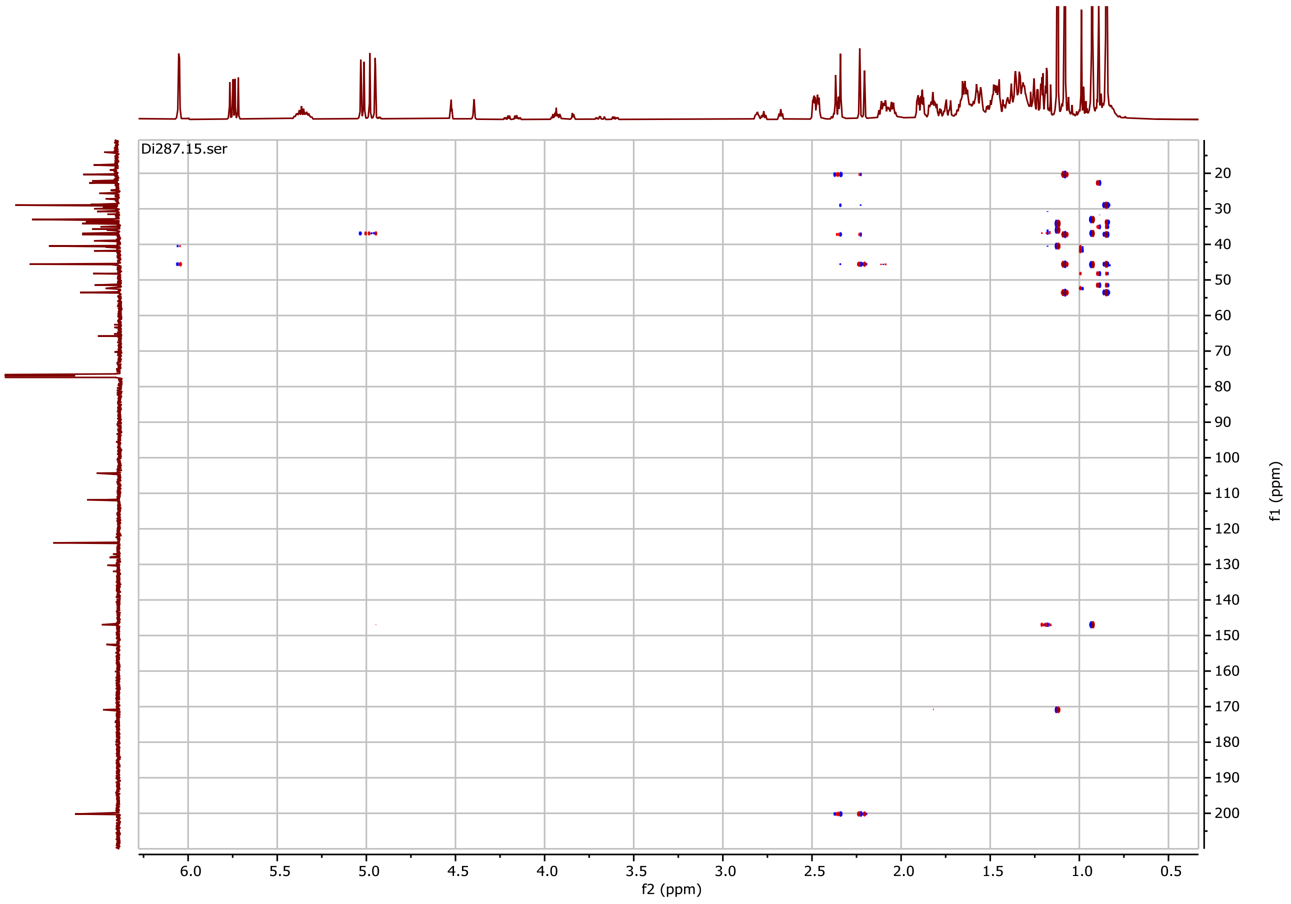


**Figure S8. gHMBCAD $^{1}H$ – $^{13}C$ NMR spectrum for the diterpenoid Di287.**

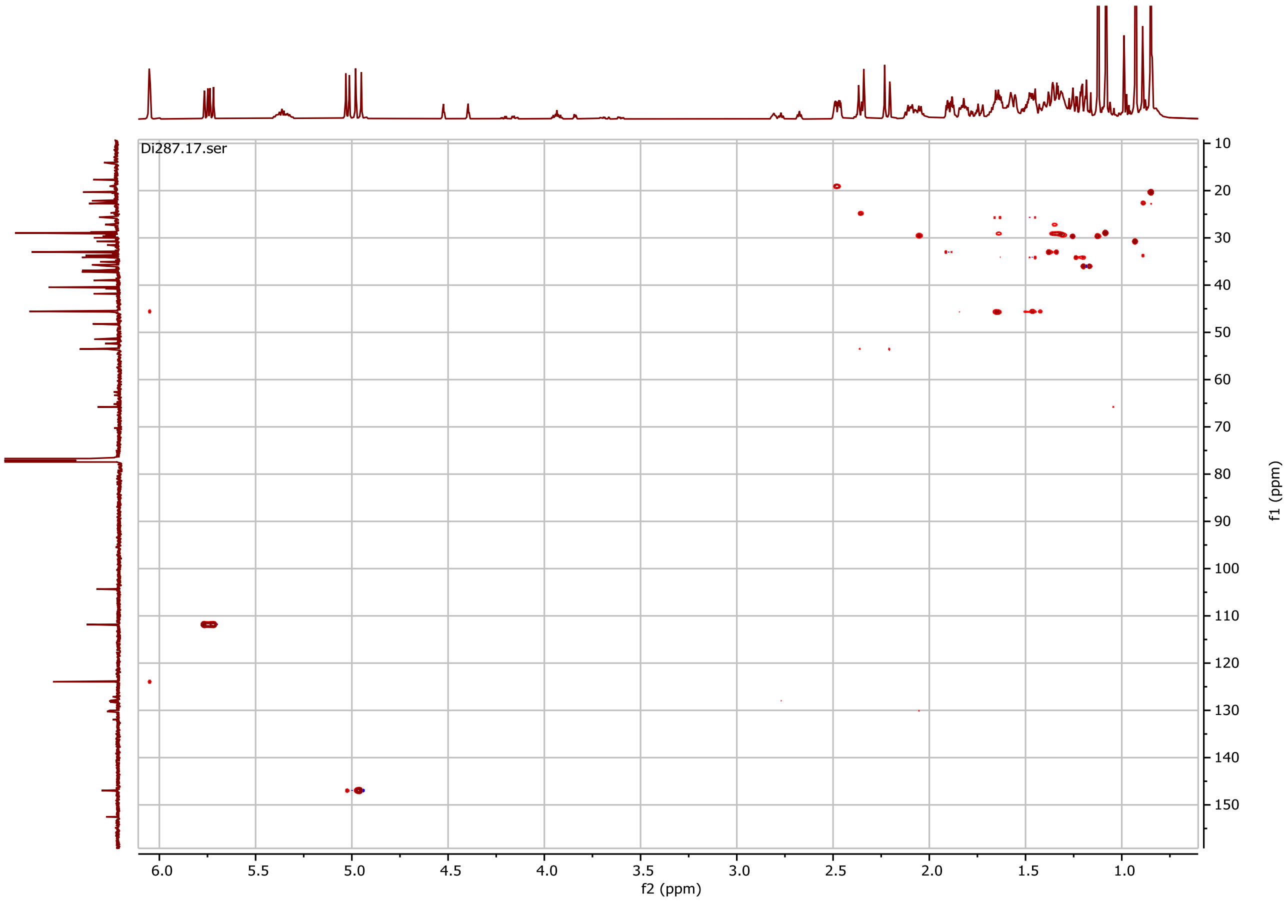


**Figure S9. gH2BCAD $^{1}H$ – $^{13}C$ NMR spectrum for the diterpenoid Di287.**

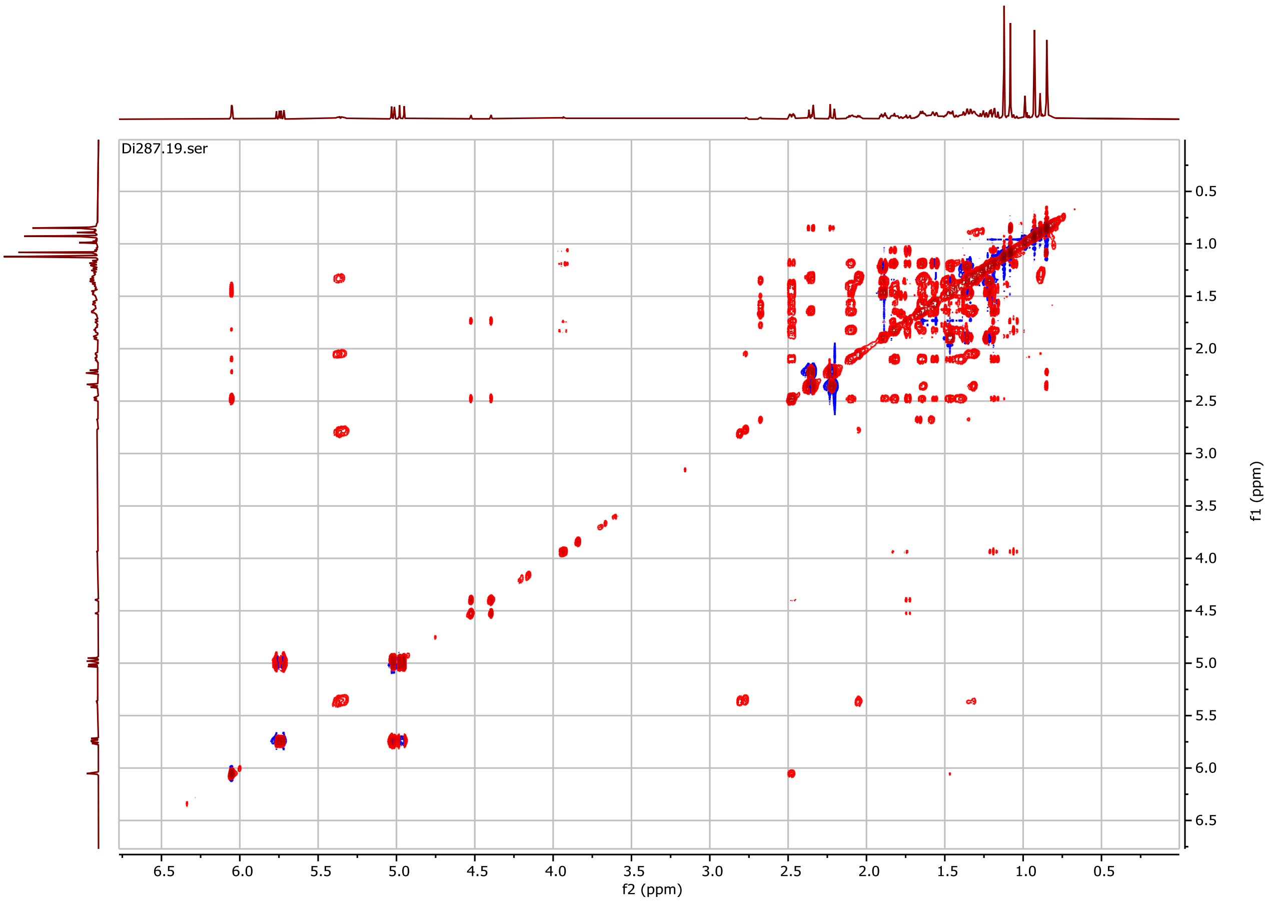


**Figure S10. gTOCSY $^{1}H$ – $^{1}H$ NMR spectrum for the diterpenoid Di287.**

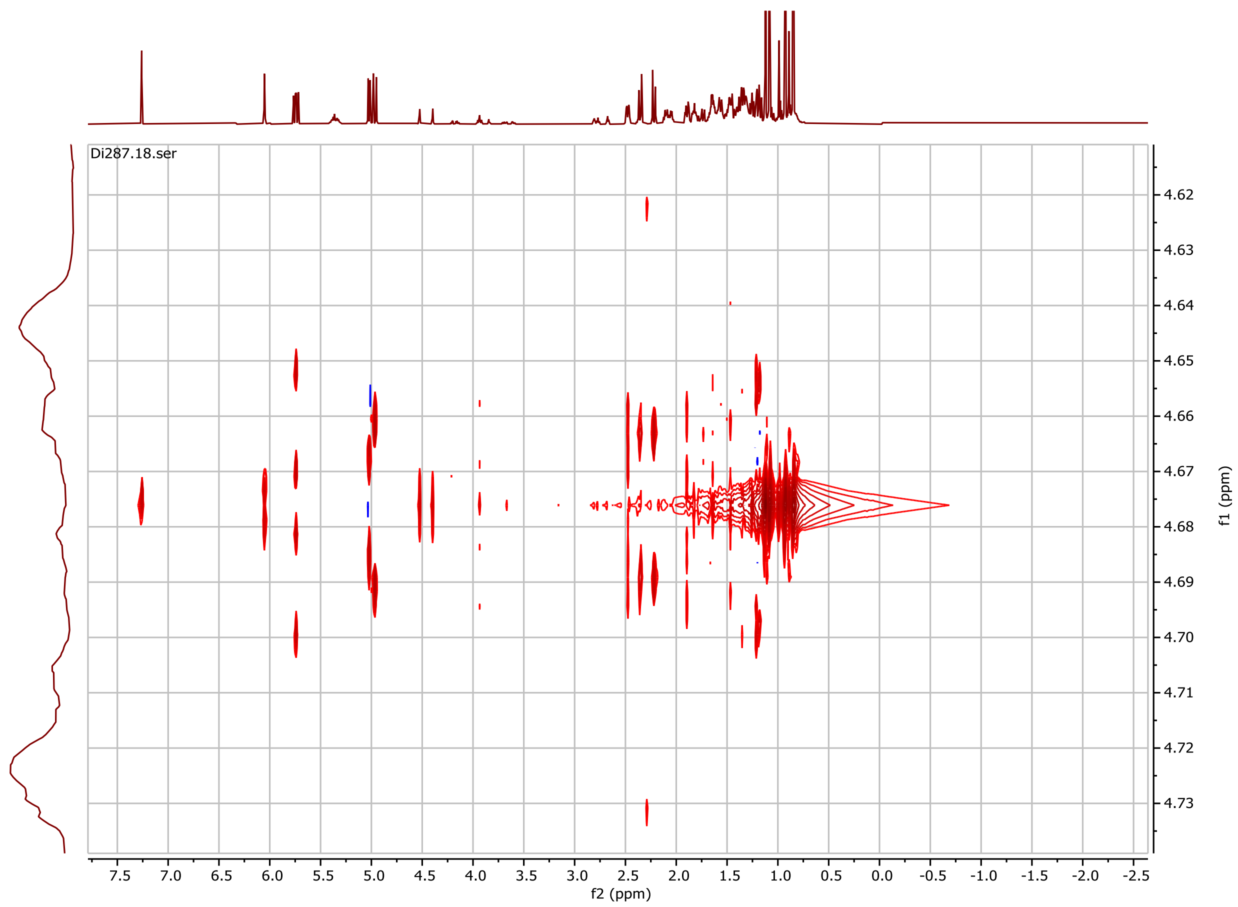


**Figure S11. HOMO2DJ NMR spectrum for the diterpenoid Di287.**

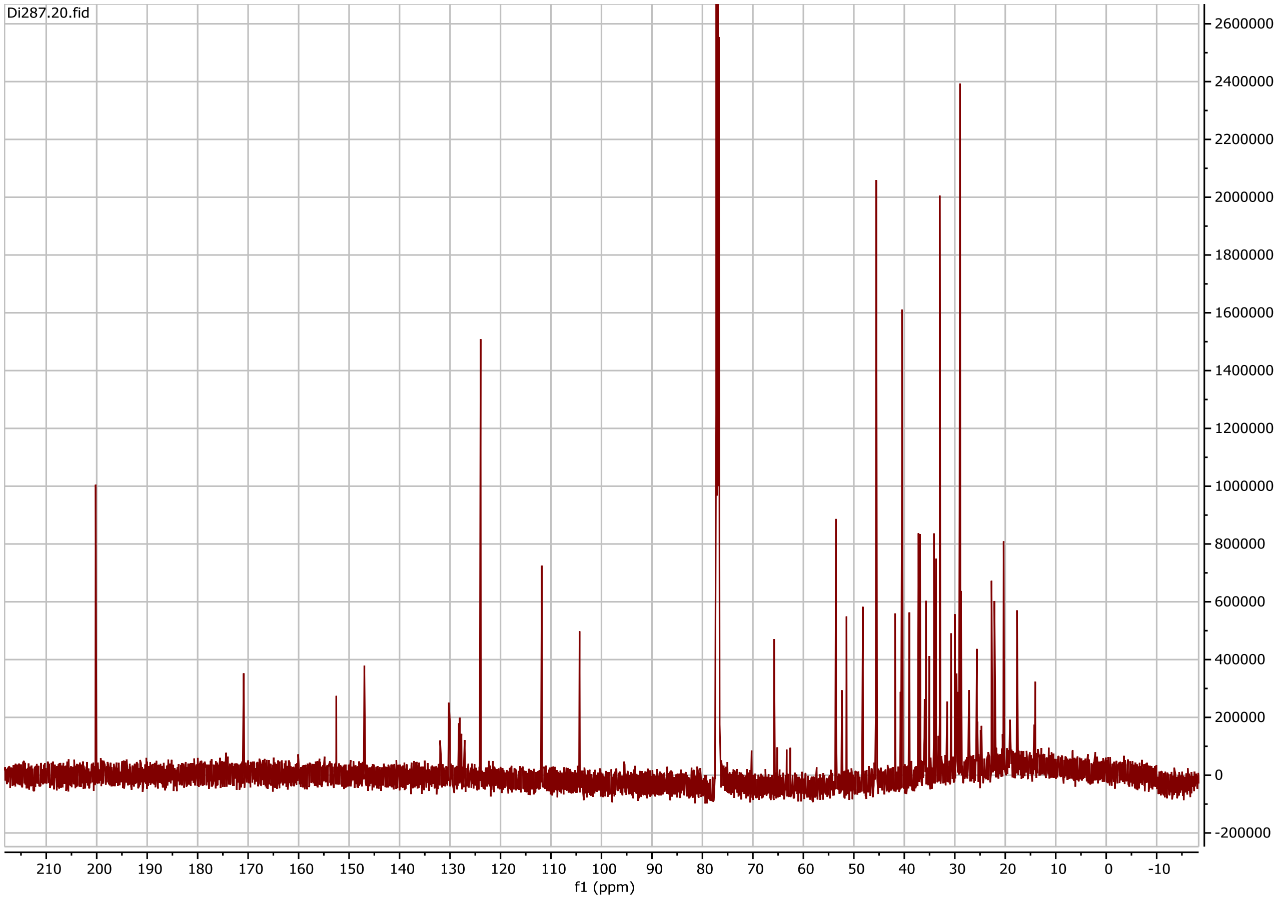


**Figure S12. $^{13}$C NMR spectrum for the diterpenoid Di287.**

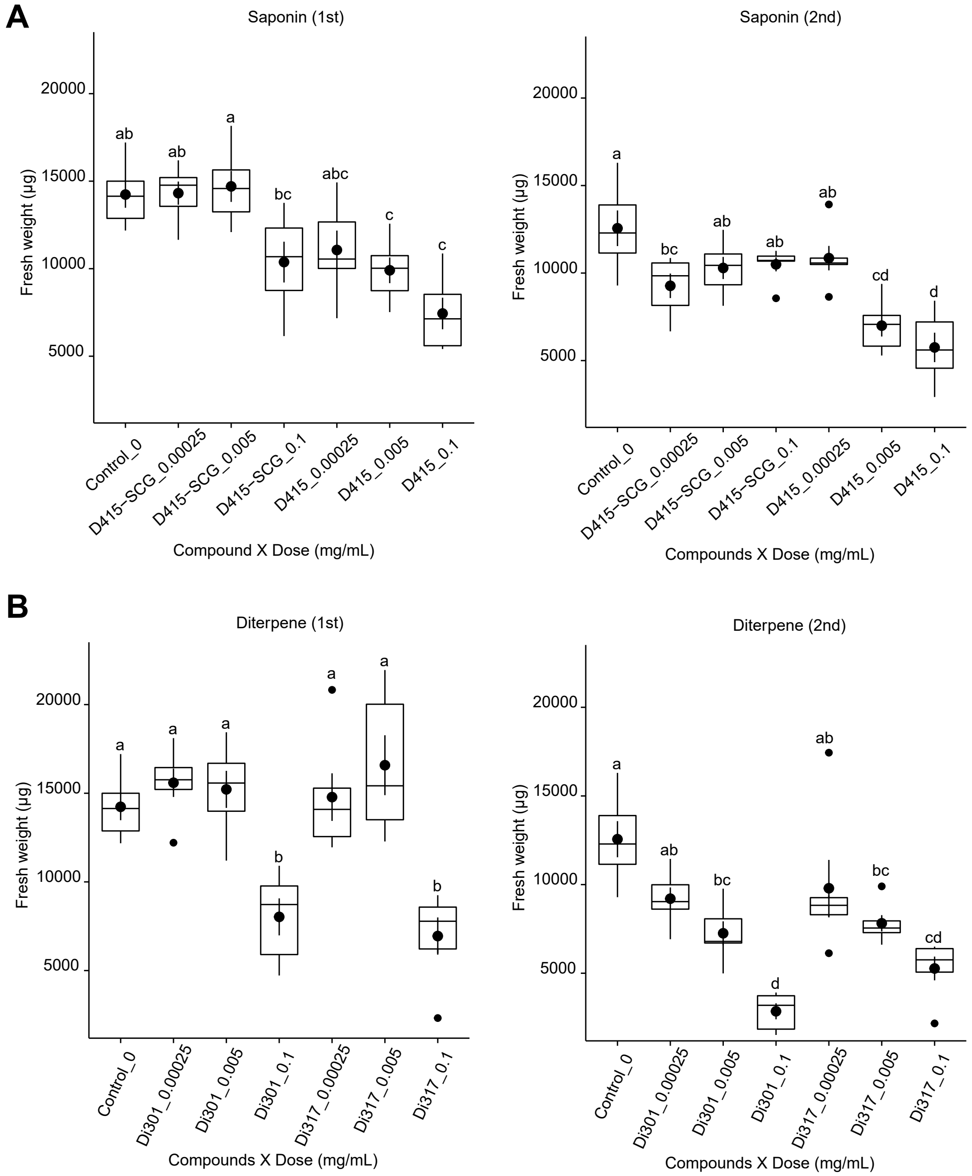


**Figure S13. The saponins and diterpenoids inhibited *Linnemannia elongata* growth in liquid-based bioassays.** Two saponins, D415-SCG and D415 (A), and two diterpenoids, Di301 and Di317 (B), were tested in two independent assays. The three tested concentrations, 0.01, 0.005 and 0.00025 mg/mL, for each metabolite treatment were used. The letters on top of the boxes indicate significant difference tested by ANOVA performed with Tukey's HSD test ($n = 6$, $FDR \leq 0.05$). $1^{st}$ and $2^{nd}$ represent independent replicates.
